# Beneficent and maleficent effects of cations on bufadienolide binding to Na^+^,K^+^-ATPase

**DOI:** 10.1101/2020.11.09.373902

**Authors:** Lucy K. Ladefoged, Birgit Schiøtt, Natalya U. Fedosova

**Affiliations:** Department of Biomedicine, Høegh-Guldbergsgade 10, 8000 Aarhus C, Denmark; Department of Chemistry, Langelandsgade 140, 8000 Aarhus C, Denmark; Natalya Fedosova. **Email:**

**Keywords:** Bufadienolide, cardiotonic steroid, ion effect, molecular dynamics, Na^+^,K^+^-ATPase

## Abstract

Kinetic properties and crystal structures of the Na^+^,K^+^-ATPase in complex with cardiotonic steroids (CTS) revealed significant differences between CTS subfamilies (Laursen et al., 2015): beneficial effects of K^+^ on bufadienolide binding strongly contrasted with K^+^/cardenolide antagonism. To solve this riddle we applied docking and molecular dynamics simulations of the complexes involving Na^+^,K^+^-ATPase, bufadienolides (bufalin, cinobufagin), and ions (K^+^, Na^+^, Mg^2+^). The results revealed that bufadienolide binding is affected by i) electrostatic attraction of the lactone ring by a cation, and ii) the ability of a cation to stabilize and “shape” the site constituted by transmembrane helices of the α-subunit (αM1-6). The latter effect was due to varying coordination patterns involving amino acid residues from helix bundles αM1-4 and αM5-10. Substituents on the steroid core of a bufadienolide add to and modify the cation effects. The above rationale is fully consistent with the ion effects on the kinetics of Na^+^,K^+^-ATPase/bufadienolide interactions.

## Introduction

The Na^+^,K^+^-ATPase, or the Na^+^-pump, is crucial for cell homeostasis and therefore represents an obvious pharmacological target. The enzyme’s abundancy, however, imposes limitations on the use of inhibitors and modulators of its activity, since they inevitably cause a number of dosedependent adverse effects and are fated to have a narrow therapeutic window. That is certainly true for cardiotonic steroids (CTS), highly specific inhibitors of the Na^+^,K^+^-ATPase, and is an issue confining their medical application. The systemic effects of an active compound may be diminished by inferring selectivity towards particular isoforms of a target protein. Thus, individual Na^+^,K^+^-ATPase isoforms revealed preferences to certain CTSs (Katz et al., 2010; Weigand et al., 2014), and these tendencies might be amplified by derivatization of the already known compounds (Katz et al., 2015). Another approach would be to design a new synthetic compound complementary to a site on the target protein. In this case, the CTS binding site is well suited as a potential point of interaction for the future drug. It has two advantages: i) it is accessible from the extracellular side, whereby negating membrane permeability as a requirement for a potential bioactive compound; ii) the site has certain plasticity since it accommodates CTSs of highly variable structures. The implementation of either strategy would gain from the availability of detailed information about the spatial organization of the binding site, modes of ligand binding, as well as ways of interfering with their interactions.

Available crystallographic data describe high affinity complexes of Na^+^,K^+^-ATPase with CTSs varying in the structure of the steroid core (cardenolides vs. bufadienolides) as well as in the degree of glycosylation (aglycones vs. mono-/triglycosylated) (Laursen et al., 2015; Laursen et al., 2013). These complexes revealed that CTSs bind to the α-subunit at the entrance to the extracellular cation transport sites. The nature of the ion bound within the site had a tremendous effect on CTS binding. Complexes with cardenolides (e.g. ouabain, digoxin) contained Mg^2+^, and substitution of Mg^2+^ with K^+^ or its congener Rb^+^ induced a re-arrangement of transmembrane helix 4 (αM4) whereby changing the position of the cardenolide in the site (Laursen et al., 2013; Ogawa et al., 2009). Functionally, it is manifested as a K^+^-induced decrease in affinity *in vitro* and described as an increased toxicity of digitalis under hypokalemic conditions *in vivo* (Fricke et al., 1981; Gelbart et al., 1980; Hansen et al., 1973). Bufalin binding, in contrast, seemed to be facilitated by K^+^ in the binding sites. The 6-membered unsaturated lactone directly coordinated K^+^ bound in cation transport site II (Laursen et al., 2015). Electrostatic pull from K^+^ and a lack of hydroxyl substituents on the steroid core allowed bufalin to slide ~1.5 Å deeper into the site. Kinetic analysis revealed that in the presence of K^+^ bufalin had higher affinity and its complex with Na^+^,K^+^-ATPase composed a homogeneous pool which allowed for crystallization. Na^+^,K^+^-ATPase/bufalin complexes formed in the absence of K^+^ had different stabilities and were clearly divided into fast- and slow dissociating pools (Laursen et al., 2015).

Thus, the above data put forward two questions: i) is K^+^-potentiation characteristic for all bufadienolides? And ii) what is the structural basis for the observed heterogeneity of Na^+^,K^+^-ATPase/bufalin complexes and how does K^+^ affect it? To find an answer, we applied an *in silico* approach (docking and molecular dynamics (MD) simulations) in conjunction with biochemical experiments. The results characterize the CTS binding cavity and ligand binding process, information that in the future will allow synthesis of compounds with predetermined properties, e.g. with improved isoform selectivity.

## Results and Discussion

The crystallized high affinity CTS complexes of the Na^+^,K^+^-ATPase are based on the E2Pi conformation of the enzyme. Therefore, all experiments in the present report describe bufadienolide’s interactions with that particular enzyme conformation exclusively, and reported values characterize equilibrium binding of CTS to the Na^+^,K^+^-ATPase in the presence of inorganic phosphate, 3 mM MgCl2, and cations of choice.

We set out by comparing the effect of K^+^ on the kinetics of Na^+^,K^+^-ATPase interactions with cinobufagin and bufalin. Cinobufagin is a bufadienolide with polar substituents on ring D of the steroid core, which might interfere with the electrostatic interactions between the lactone ring and K^+^ in the cation binding site described for bufalin.

Figure 1 compares the residual activity of Na^+^,K^+^-ATPase after enzyme pre-incubation with varying concentrations of bufalin and cinobufagin in the presence or absence of K^+^. The degree of inhibition corresponds to an amount of the enzyme-inhibitor complex, and kinetic analysis of these curves allows for the extraction of apparent affinity values for each inhibitor (Laursen et al., 2015). Both bufadienolides exhibit high affinities toward Na^+^,K^+^-ATPase in the absence of any other ions but Mg^2+^. As shown previously, bufalin affinity was high (Kd = 0.01 μM), yet, incomplete inactivation revealed formation of both fast- and slow-dissociating complexes (4). Presence of K^+^ improved bufalin affinity and prevented formation of the fast-dissociating complex (Figs. 1 and S1, Table S1). Monitoring of bufalin substitution with anthroylouabain estimates the dissociation rate constant in the absence of K^+^ as 5.7·10^-4^ s^-1^. It becomes 300 times slower in the presence of K^+^ (2.2·10^-6^ s^-1^, Fig. 1B). Note, that the koff from the fast-dissociating complex could not be determined in our experimental setup. This clear K^+^-stabilizing effect on the Na^+^,K^+^-ATPase/bufalin complex was only partially observed for cinobufagin. Cinobufagin binding in the absence of K^+^ revealed the same pattern as that of bufalin: high affinity (K_d_ = 0.05 μM) but incomplete inhibition (Table S1). The proportion between fast- and slow-dissociating complexes was essentially identical to that of bufalin. Addition of K^+^ promoted inhibition, i.e. eliminated the formation of the fast-dissociating complex, but the affinity for cinobufagin decreased (Fig. 1A). The latter resembles the K^+^ effect on cardenolide binding to Na^+^,K^+^-ATPase. Thus, the consequences of K^+^ presence for bufalin and cinobufagin binding are not alike. While K^+^ ensures homogeneity of both bufadienolide/enzyme complexes, it has opposite effects on their apparent affinities.

**Figure 1.**
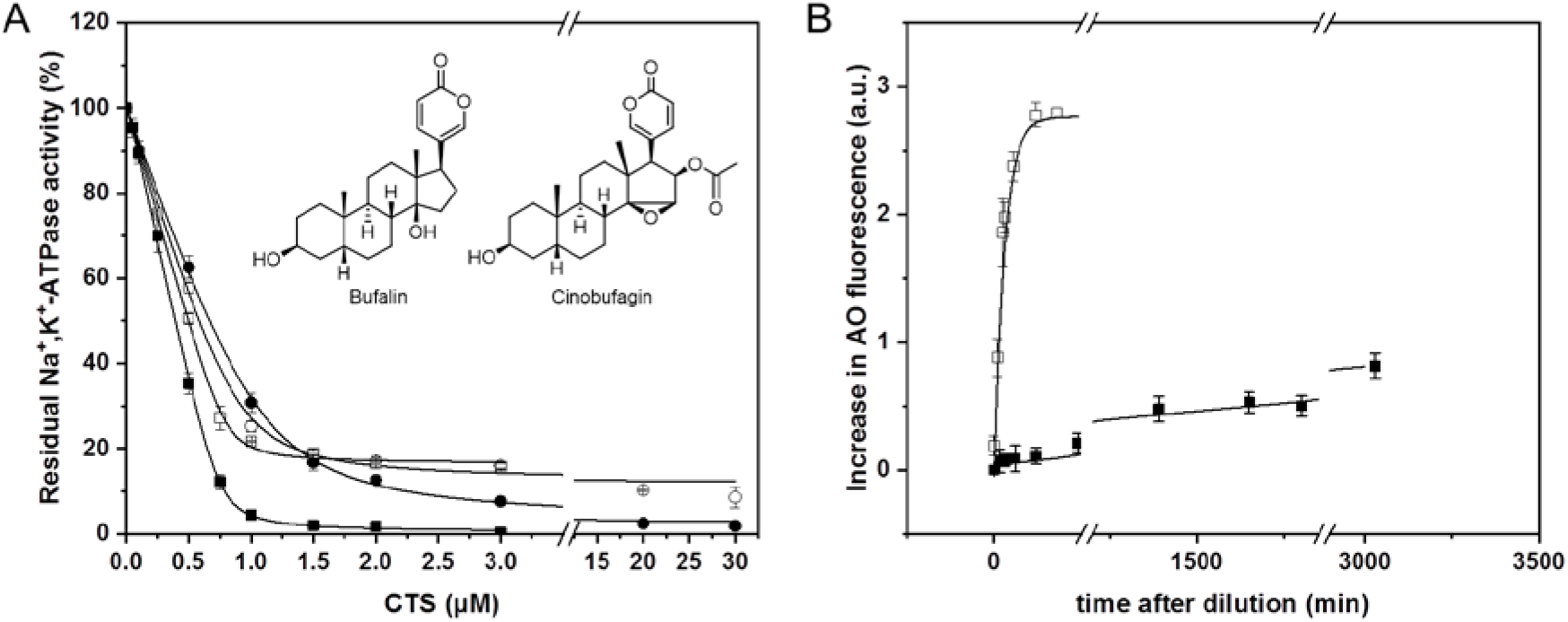
Interactions of bufadienolides with Na^+^,K^+^-ATPase. (A) Binding of bufalin (squares) and cinobufagin (circles) in the presence or absence of K^+^ (filled and open symbols, respectively). The amount of CTS-bound enzyme was estimated from percentage of inhibition of the enzyme activity. The data were fitted to a square-root equation as described in ref. (Laursen et al., 2015), the obtained parameters are collected in Table S1. (B) Time course of bufalin dissociation from the complex with Na^+^,K^+^-ATPase in the presence (filled squares) or absence (open squares) of 10 mM KCl. Bufalin dissociation was induced by dilution and addition of anthroylouabain (AO). The dissociation rate constant obtained from the monoexponential fit of the data was 2.2·10^-6^ s^-1^ in the presence of K^+^ and increased to 5.7·10^-4^ s^-1^ in the absence of K^+^. All data points represent the average of at least three experiments.

In order to understand the mechanisms underlying the multiple and diverse K^+^ effects on bufadienolide binding to Na^+^,K^+^-ATPase we turned to computational experiments. It is not feasible to obtain crystallographic evidence of the complexes of interest due to their transient nature. First, we performed docking calculations to probe the possible binding modes of each bufadienolide in the CTS binding site. The docking calculations consider both ligand and protein flexibility which allows for more variation in the binding modes. They were performed for bufalin and cinobufagin in Na^+^,K^+^-ATPase with and without two K^+^ ions bound to ion site I and II. The resulting poses of each bufadienolide were clustered according to their conformational and positional similarity, and the output of the calculations is summarized in Table S2.

Bufalin always bound with the lactone ring facing down; an upside down orientation of that compound (as previously suggested for an ouabain derivative (Sandtner et al., 2011)) was never observed. Its docking into Na^+^,K^+^-ATPase with K^+^ ions in place resulted in three binding clusters (Fig. S2). The first, denoted B4_K+_, shows bufalin bound very close to the extracellular surface in a near vertical position, while in B5_K+_ and B6_K+_ the ligand has an overall position similar to that in crystal structures (Laursen et al., 2015). In B5_K+_, the methyl substituents on the convexβ surface of bufalin point toward αM4, while in B6_K+_ they point toward αM2. The majority of the docking output belongs to the B6_K+_ cluster in agreement with the crystal structure of the Na^+^,K^+^-ATPase/bufalin complex. The same three binding clusters, although now almost equally populated, were found in docking to the site in the absence of K^+^ (Table S2). Interestingly, the rare, vertical binding mode denoted B4_k+_ becomes common in the absence of K^+^; however, such changes are difficult to interpret based solely on a docking calculation.

Docking of cinobufagin in the presence of K^+^ resulted in only one binding cluster, denoted C3_k+_, which was highly similar to the crystallized binding mode of bufalin in Na^+^,K^+^-ATPase (Laursen et al., 2015) and thus also cluster B6_K+_ of bufalin (Figs. S2 and S3). In contrast, the number of binding clusters increased to five in the absence of K^+^ (denoted C1, C3, C4, C5, and C6 on Fig. S3). C1, which includes the highest number of poses, is similar to the crystallized binding mode of the ligand in the Na^+^,K^+^-ATPase/bufalin complex (Laursen et al., 2015), and thus also C3_K+_ and B6_K+_, while C3 is a vertical binding mode similar to what was observed for bufalin above in B4_K+_. C4-C6 all represent upside down binding modes in which the lactone ring points out of the binding site.

Collectively, the docking calculations suggest that the positive charge of K^+^ in cation binding site II assists in proper orientation of the bufadienolides and even coordinates directly to the 6membered unsaturated lactone. Steric clashes from the substituents on the cinobufagin steroid core, however, may be the reason for the decreased affinity despite the stabilizing electrostatic impact.

Thus, K^+^ in site II is important for bufadienolide binding due to its electrostatic interactions with the lactone ring. Is this property specific for K^+^ or will any cation in the binding site exert the same navigating/stabilizing effects? Crystal structures have revealed that the extracellular cation sites accept other ligands than K^+^ and its congeners, e.g. Mg^2+^ and Mn^2+^ (Laursen et al., 2013). We therefore set out to test the effect of Na^+^ and Mg^2+^ on bufalin binding.

Figure S1 and Table S1 summarize the effects of Mg^2+^, K^+^, and Na^+^ in various concentrations on the apparent affinities of bufalin. It is clear that only K^+^ prevents formation of the fast-dissociating complexes and improves affinity. The effect is effectuated at very low K^+^ concentrations as expected, as the affinity of the extracellular binding sites for K^+^ is high (Fig. S1A). An increase in Mg^2+^ concentration (above 3 mM Mg^2+^ i.e. standard condition) has no further effect on bufalin binding.

The sites are therefore either saturated with Mg^2+^ under standard conditions, or Mg^2+^ does not affect bufalin binding (Fig. S1B). Na^+^-affinity for the extracellular sites, on the other hand, is very low since the change in bufalin binding kinetics demands a high Na^+^-concentration (Fig. S1C). Importantly, the effects of Na^+^ and K^+^ are ion-specific since 200 mM NMG^+^ (N-methyl-D-glucamine), used as a control for the effect of ionic strength alone, had no influence on bufalin binding (Fig. S1B).

Thus, the effects of K^+^ on bufadienolides’ interactions with the Na^+^,K^+^-ATPase are distinct from that of other ions bound to the extracellular sites. This finding triggered MD simulations of the Na^+^,K^+^-ATPase/bufadienolide complexes in the presence of different ions. Analyses of the dynamical aspects of bufadienolide binding aims to reveal the structural consequences of the ions present in the binding site.

MD simulations were performed with the crystallized Na^+^,K^+^-ATPase/bufalin complex (Laursen et al., 2015) as a starting structure in the presence of the ions tested above as well as an apo conformation. Simulations with K^+^ considered full occupation of sites (i.e. two K^+^ ions). Since Mg^2+^ is mandatory for phosphorylation and always present in the biochemical experiments, the series included Mg^2+^-bound enzyme. The Na^+^ bound system was simulated with both one and two Na^+^ ions since affinity for Na^+^ was low. Singular ions were always located in site II. The Na^+^,K^+^-ATPase/cinobufagin complex obtained from the docking calculation (C3_K+_) was simulated with either two K^+^ ions or one Mg^2+^ ion bound. Each system was simulated for 500 ns in three repeats, except for the Na^+^,K^+^-ATPase/bufalin/2xK^+^ system which was simulated in five repeats (MD1-5). Each molecular system was evaluated for: *i)* ligand stability within the binding site, *ii)* ligand/ion coordination (site II), *iii)* ion/protein coordination (site II), and *iv)* protein/ligand interaction strength. Combined, these four metrics characterize both the direct ion effect through coordination of the ligand and the indirect ion effect due to stabilizing the binding site around the bufadienolide.

A caveat of this study is the physical description of ions by the force field. In the force field applied in this work, as well as in the majority of biologically relevant force fields, each ion is represented by a point charge and two parameters describing the van der Waal forces. By construction, the description is symmetric, and the ions are distinguishable by the coordination distance alone and not by the coordination geometry. As many details are lost, the ions, especially the divalent ions, behave too similarly in MD simulations (Yu et al., 2010). Nevertheless, focusing on the reproducible aspects, such as the coordination distance, we obtain reliable data.

A clustering calculation provided an initial overview of the behavior of the bufadienolides within the binding site. In all cases the steroid core was mainly stabilized by hydrophobic interactions and the primary difference was in the rotational state of the bufadienolide. Across all systems, four major binding modes (BM0-3) were determined (Figure 2). Thus, in BM0 the β-surface of the bufadienolide points toward αM2 in agreement with the crystal structure of the Na^+^,K^+^-ATPase/bufalin complex (Laursen et al., 2015). In BM1, BM2, and BM3, the β-surface points toward αM1, αM4, and αM6, respectively. The core hydroxyl group (C14_β_) of bufalin forms a hydrogen bond with Thr797 only in BM0 and BM1. An equivalent hydrogen bond was observed in crystal structures for all CTS-enzyme complexes (Gregersen et al., 2016; Laursen et al., 2015; Yatime et al., 2011). The equivalent epoxy group on cinobufagin is not able to form a hydrogen bond to Thr797 due to the altered local conformation near the oxygen atom leading to a suboptimal hydrogen bonding angle. In addition to the rotational state, the depth of binding, i.e. z-coordinate, showed some degree of variation. Thus, in order to assess stability of the bufadienolides within the site it is necessary to track their rotational state as well as the depth of their location. Figure 3 presents an overview of ligand stability for all systems. Bufalin bound to Na^+^,K^+^-ATPase in the presence of K^+^ is stable and in BM1 (β-surface facing αM1) despite being initiated from BM0. Bufalin changes from BM0 to BM1 within the first ns. Minor fluctuations in binding depth were detected in all repeats except MD2 and the last 100 ns of MD4 where it moves even further down the binding site. In the presence of one or two Na^+^ ions the behavior of bufalin is highly similar. In simulations with Mg^2+^, bufalin is less stable and retains BM1 only in one repeat simulation. Instead, bufalin binds in all four major binding modes and in additional minor binding modes in varying depths. The movements up and down the site have larger amplitudes than otherwise observed except for the apo conformation, where bufalin retains BM1 exclusively, but moves up and down in the binding site even more. In fact, in two simulations (MD1 and MD2) bufalin almost exits to the extracellular milieu (Fig. S4). Intriguingly, the tilted binding mode observed in the docking calculations reappeared in MD1 as a semi-stable binding mode during bufalin’s exit path (Fig. S4B).

**Figure 2.**
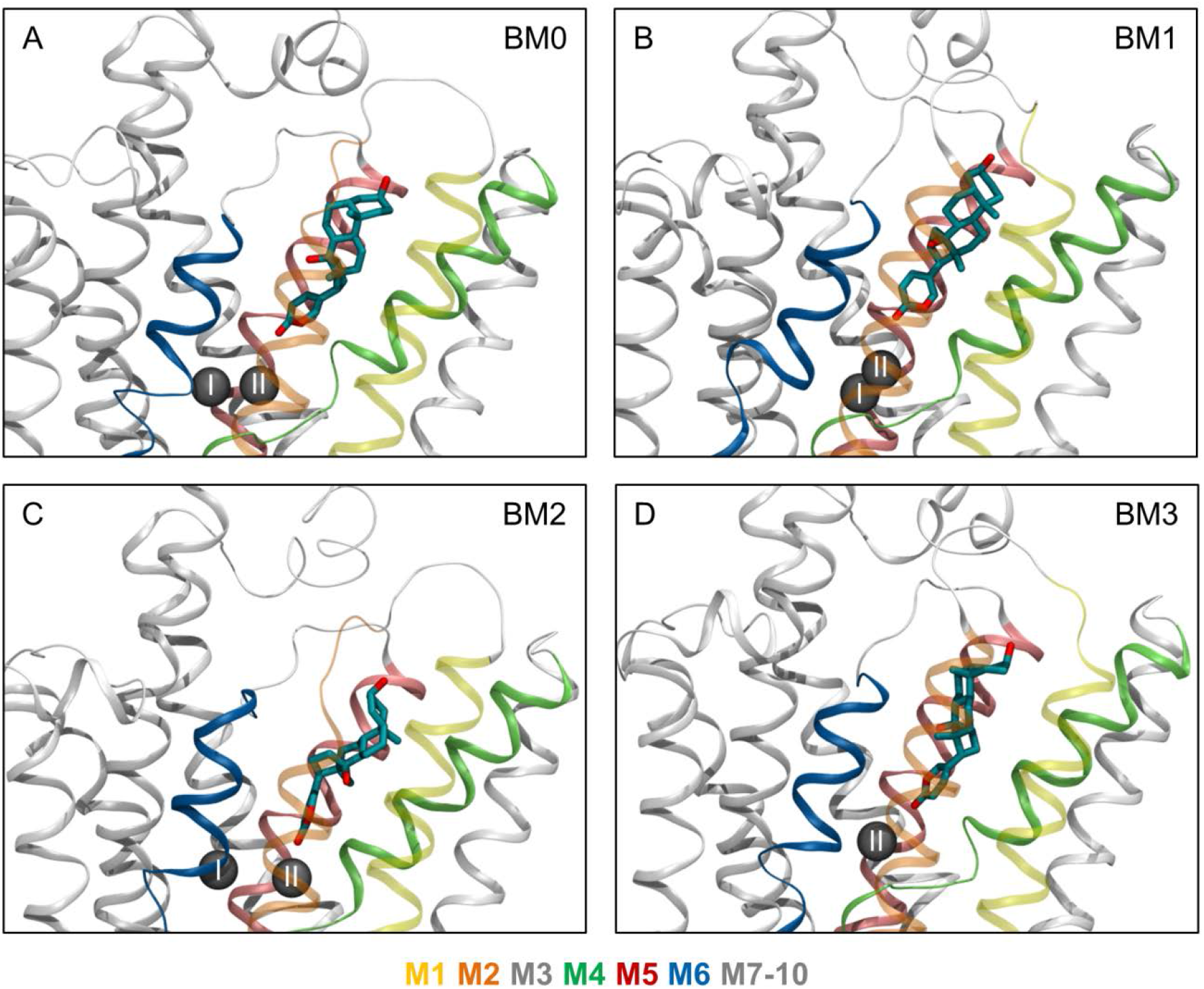
Common binding modes of bufalin in Na^+^,K^+^-ATPase. **(A)** The binding mode observed by X-ray crystallography (PDB ID: 4RES) denoted BM0. The three most common binding modes observed in MD simulations denoted **(B)** BM1, **(C)** BM2, and **(D)** BM3. Each binding mode corresponds to a specific rotational state of bufalin within the enzyme: in BM0 the β-face of bufalin points toward αM2, in BM1 it points toward αM1, in BM2 toward αM4, and in BM3 toward αM5. Bufalin is shown in cyan, K^+^ ions as gray spheres, and the enzyme as ribbons. αM1 and αM2 are shown in transparent ribbons to not obstruct the view of bufalin.

**Figure 3.**
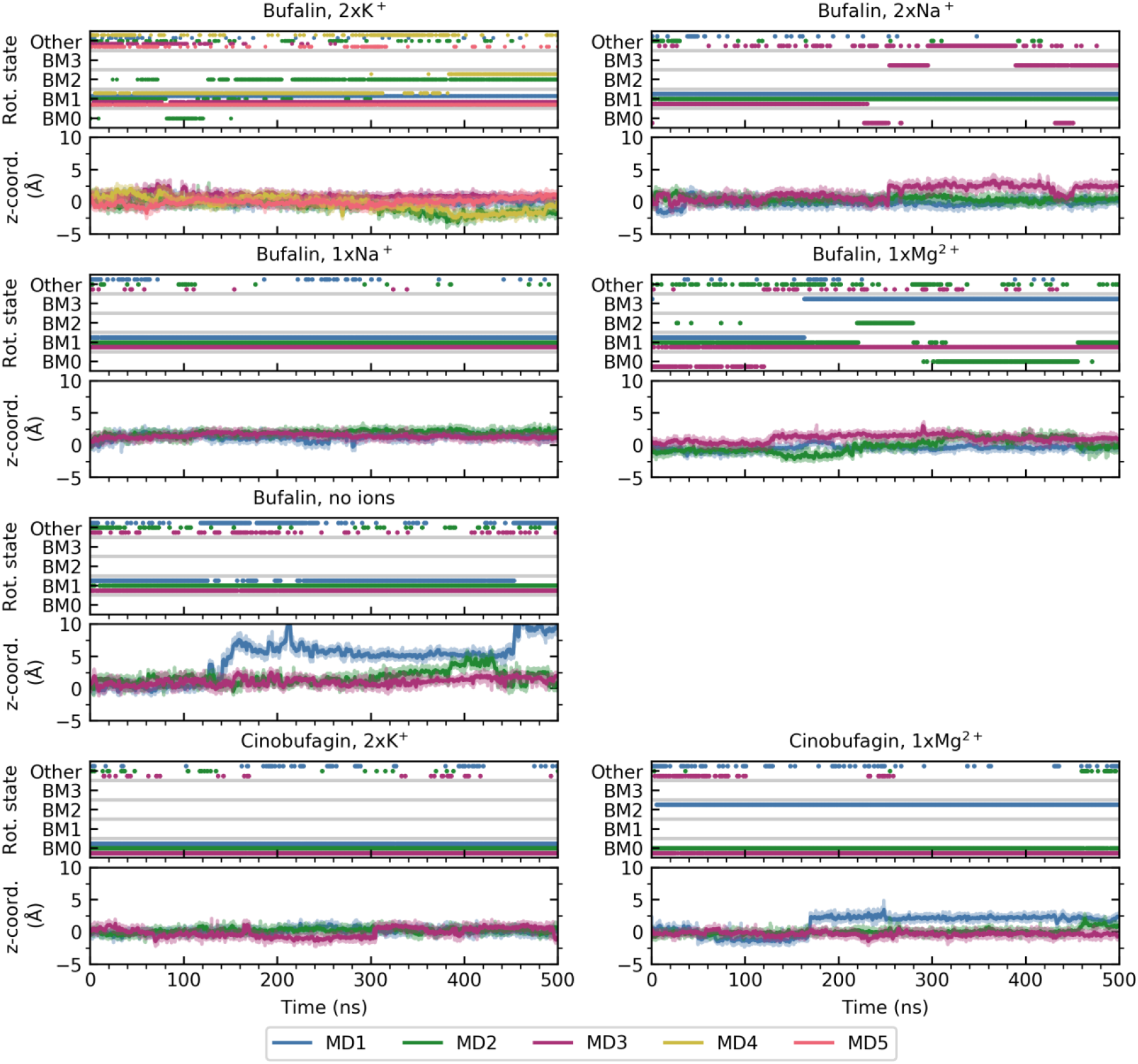
Bufalin and cinobufagin stability in the binding site in the presence of Na^+^, K^+^ and Mg^2+^. The stability is evaluated from the rotational state of the bufadienolide (BM0-3), and for each rotational state from the fluctuations up and down the binding site reflected in the z-coordinate of the bufadienolide center of mass relative to the starting structure. The rotational state was obtained from the conformational clustering of the data. All trajectories were aligned prior to analysis using αM5-10 of the enzyme. Simulation repeats are named MD1-5 and colored separately.

Overall, BM1 appears to be a preferable binding mode of bufalin as it appeared in the majority of the simulation repeats regardless of ions bound, with the exception of Mg^2+^. The crystal structure of the Na^+^,K^+^-ATPase/bufalin complex, however, represents BM0 (Laursen et al., 2015) with the steroid core rotated almost 45° compared to BM1. This rotation alters the position of the hydrophobic β-surface in such a way that it faces the apolar αM1 (BM1) instead of the considerably more polar αM2 (BM0). The difference between the MD simulation results and the crystallized form with a less favorable orientation of bufalin might be caused by the crystal packing. Note, that all CTS crystallized in complexes with Na,K-ATPase have the same binding mode (BM0) irrespectively of the hydrophobicity of their steroid core (Gregersen et al., 2016; Laursen et al., 2015; Yatime et al., 2011).

The Na^+^,K^+^-ATPase/cinobufagin complex was simulated in the presence of either K^+^ or Mg^2+^ ions. With K^+^ in the sites, cinobufagin was overall stable in BM0 (β-surface facing αM2) albeit positioned slightly higher up in the binding site compared to bufalin (Fig. 3). Closer inspection of the simulations revealed that this mode was best at accommodating the larger acetyl group of the core above the shorter αM6 helix. With Mg^2+^ in the site, BM0 was observed in two repeats, while BM2 was observed in one (MD1). Cinobufagin in BM2 moves a few Ångström further out of the site (toward the extracellular milieu). Thus, cinobufagin is most stable in the presence of K^+^ ions, although the large acetyl group prevents optimal coordination with K^+^ in site II. The preferable binding mode of cinobufagin, BM0, is consistent with crystal structures of other CTS (Gregersen et al., 2016; Laursen et al., 2015; Yatime et al., 2011).

Monitoring of the coordination distances for the ion in site II is a mode of evaluation of its direct interaction with the lactone oxygen of the bound bufadienolide. The first-shell coordination distance of Mg^2+^, Na^+^, and K^+^ is approximately 2.1 Å, 2.4 Å, and 2.8 Å, respectively (Zheng et al., 2008). In the calculations, direct coordination was defined as distances shorter than the optimal coordination distance plus 0.1 Å.

Simulations of the Na^+^,K^+^-ATPase/bufalin/2xK^+^ system revealed that distribution peaks at 2.8 Å were most populated (Figure 4). They indicate direct coordination and were maintained in 63 % of the total simulation time of MD2-MD5. MD1 was an exception: the direct coordination occurred only in < 2 % of the time. Instead, we registered a peak at 4.5 Å. Thus, the distance distributions suggest that K^+^ does not always interact with bufalin, but direct coordination is common.

**Figure 4.**
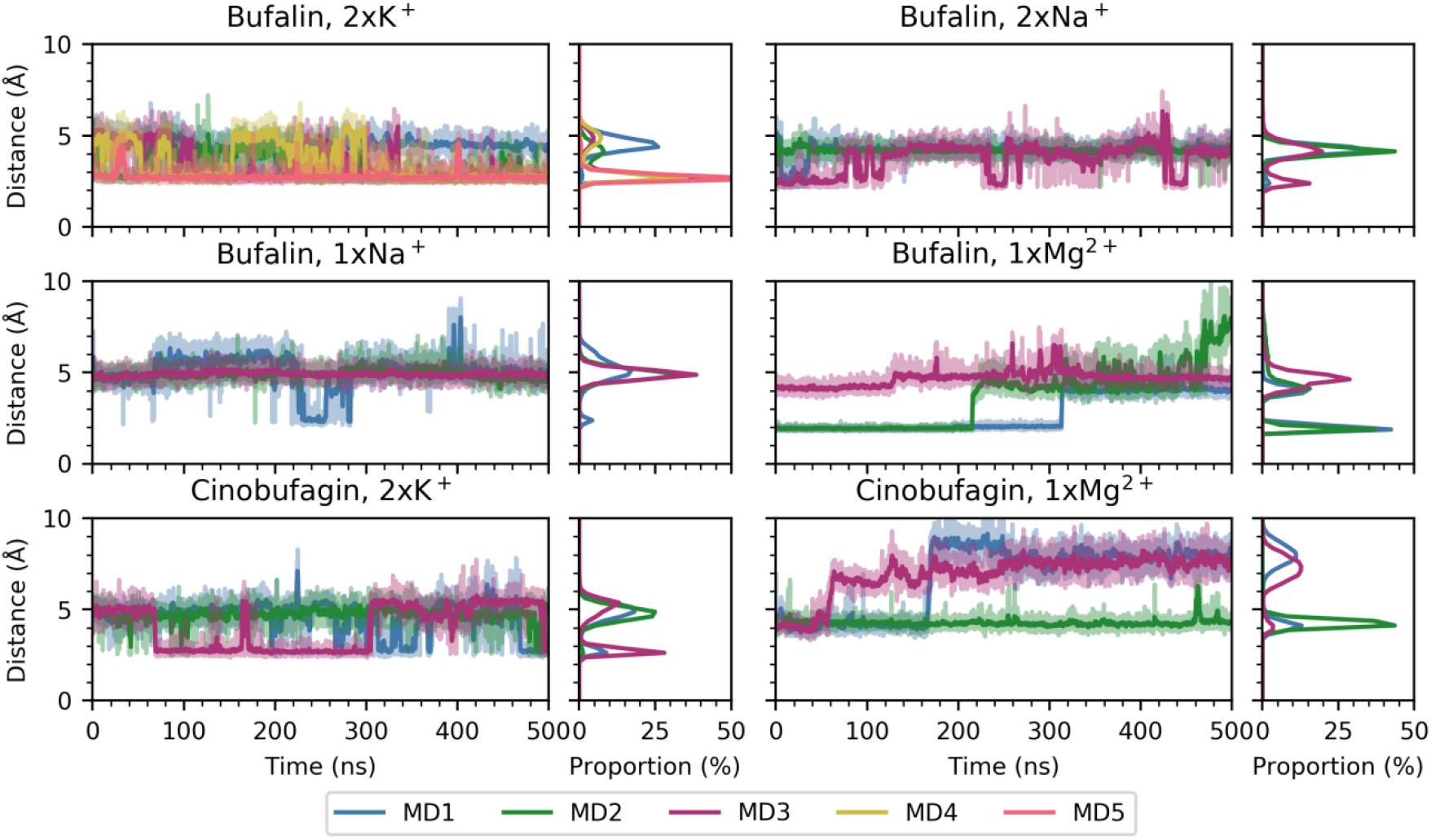
Direct coordination of the ion in cation site II by bufalin and cinobufagin. The distance between the ion in site II and carbonyl oxygen in the bufadienolide lactone ring as a function of time is reported for each molecular system. The running average is shown in opaque hues while the raw data is shown in transparent hues based on the color legend below the plot. The resulting distance distribution (not fitted) is shown on the right hand side of each plot.

Simulations of Na^+^,K^+^-ATPase/bufalin/2xNa^+^ revealed two peaks at 2.4 and 4.2 Å. In all three repeats, the second (distant) peak was most populated, while direct coordination of the ion was only detected in ~7 % of the total simulation time. The same trend, with even fewer occurrences of direct coordination (<2 %), was observed with a single Na^+^ present.

Analysis of the Mg^2+^-simulations revealed two peaks, but the populations were not similar across repeats. In MD1 and MD2, direct coordination was observed in approximately half of the simulation time (62 and 43 %, respectively), while in MD3, direct coordination was never observed. Simulations revealed “irreversibility” of the breakage of direct coordination: when the direct coordination was broken, the interaction distance remained long for the rest of the simulation time. This feature is in contrast to the other simulations, where direct coordination broke and restored multiple times. Visual inspection revealed that coordination broke as Mg^2+^ tended to move deeper into site II, while bufalin was unable to follow Mg^2+^ due to steric hindrance by the enzyme.

Simulations of Na^+^,K^+^-ATPase/cinobufagin/2xK^+^ also displayed two distance distributions, but direct coordination to cinobufagin was only observed in 19 % of the combined simulation time, in sharp contrast to bufalin. Direct coordination broke and re-formed multiple times. In simulations of Na^+^,K^+^-ATPase/cinobufagin/Mg^2+^, direct Mg^2+^-coordination was observed initially in all three repeats, but only consistently in MD2. The other repeats were similar to the Na^+^,K^+^-ATPase/bufalin/Mg^2+^ simulations where direct coordination was never restored after initial loss.

In summary, bufalin in the preferable BM1 binding mode, maintained direct coordination of K^+^ in site II most of the simulation time. In contrast, the BM1 mode of bufalin is not reconcilable with Na^+^ and Mg^2+^ binding, and direct Na^+^ and Mg^2+^ coordination by bufalin was rare. In both cases, bufalin can either move deeper into the site and achieve direct coordination, or move higher up the site and occupy the second coordination shell of the ion (Varma et al., 2006). In practice, the latter was detected for bufalin, which would result in decreased bufalin affinity. Cinobufagin favored the BM0 binding mode, likely due to steric hindrances from its large acetyl moiety, and direct coordination by both K^+^ or Mg^2+^ was rare.

The paragraph above describes direct interactions of the ion in site II with the bufadienolide of choice. The ion, however, also coordinates the protein and may thus alter the geometry of the CTS binding site wedged between the flexible αM1-4 helices and the rigid αM5-M10 bundle. In the simulations, the ions in site II are coordinated by the backbone carbonyl of Val325 (αM4) and one or two of the side chain oxygen atoms of Glu327 (αM4), Glu779 (αM5), and/or Asp804 (αM6) (Fig. 5A). Coordination to Ala323 was not detected (Laursen et al., 2015; Ogawa et al., 2009). Since it is feasible that coordination of an ion by both helix bundles simultaneously stabilizes the composite CTS site, the duration of coordination in conjunction with the helical location of a given coordinating residue may serve to quantify the indirect influence of an ion on CTS binding. It is particularly relevant for bufalin and cinobufagin as their β-surface has fewer polar substituents than most CTS.

**Figure 5.**
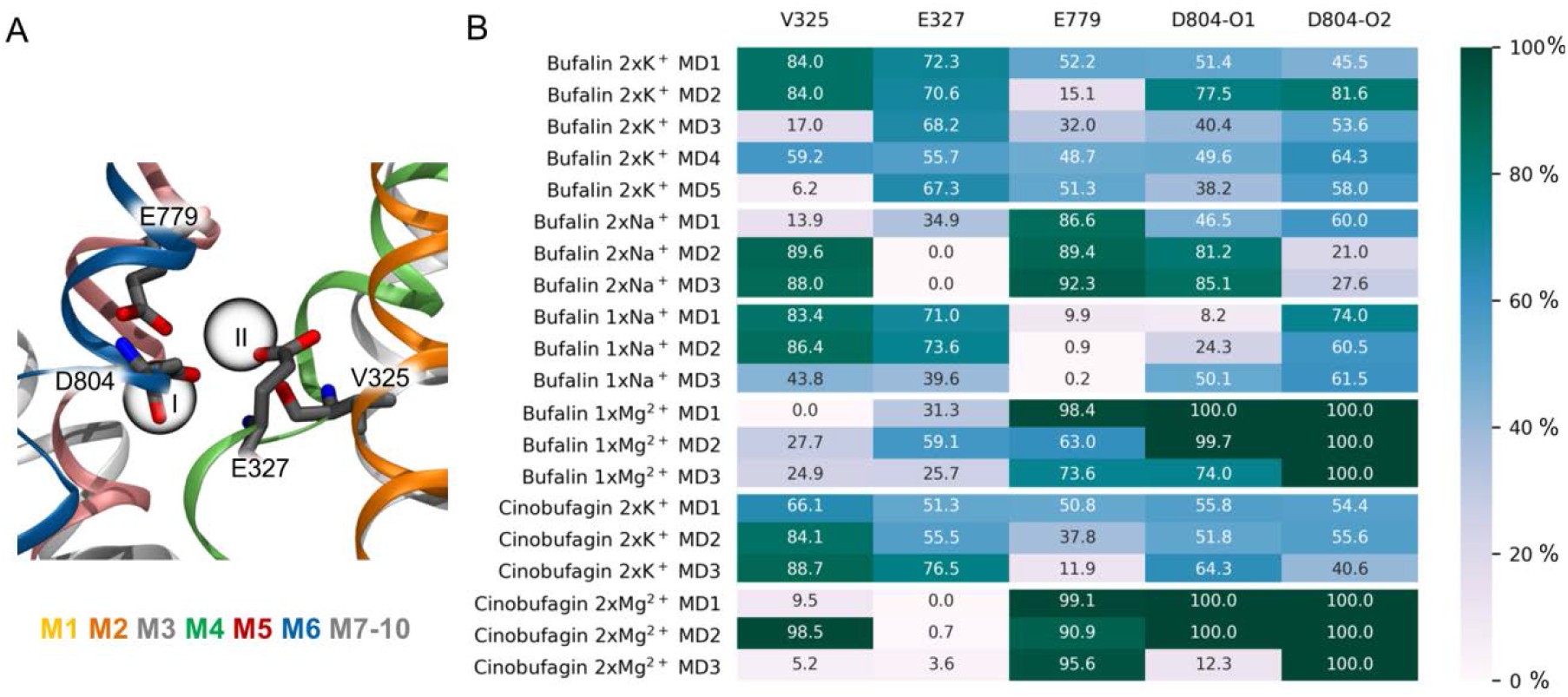
Prevalence of ion (site II) coordination to residues within Na^+^,K^+^-ATPase. (**A)** Structural representation of ion site I and II (gray spheres) and the residues that can coordinate the ion in site II; namely the backbone oxygen atom of Val325 (αM4), one side chain oxygen atom from Glu327 (αM4), one side chain oxygen atom from Glu779 (αM5), and either side chain oxygen atom from Asp804 (αM6). (**B)** Duration of direct ion coordination expressed in per cent of total simulation time. Coordination was considered as direct when the distance was shorter than the optimal coordination distance plus 0.1 Å (i.e. 2.2 Å (Mg^2+^), 2.5 Å (Na^+^), and 2.9 Å (K^+^). Each cell is colored according to the color scale bar right of the plot with high percentages being dark green and low percentages being light gray.

In the Na^+^,K^+^-ATPase/bufalin/2xK^+^ simulations, we observed coordination of K^+^ by Glu327 and Asp804 (single and double coordination, respectively) in approximately 50-75 % of the simulation time (Fig. 5B). Coordination to Val325 varied more and ranged from 6 to 84 % depending on the repeat simulation. Some coordination to Glu779 in up to half of the simulation time was also present. The overall coordination pattern observed in double Na^+^ occupation was very similar to that in double K^+^: Na^+^ was consistently coordinated by Val325, Glu327, and Asp804. In contrast to K^+^, Na^+^ did not coordinate to Glu779, and double coordination by Asp804 was rare. However, since Na^+^ affinity to E2P is low, the existence of the E2P(2Na)/bufalin form might be irrelevant under physiological conditions. Coordination of a single Na^+^ ion was different: it was very strong to Glu779, and coordination by Val325 and Asp804 were also common. However, coordination to Glu327 and double coordination by Asp804 were rare. This altered behavior could be a consequence of a loss of stabilizing effect of the ion in site I on the ion in site II. The RMSD of the ion in site II was sometimes higher when only one site was occupied (Fig. S5). The single Mg^2+^ ion in site II was strongly coordinated to Glu779 and doubly coordinated to Asp804, while its coordination to Val325 and Glu327 were rarer. The Na^+^,K^+^-ATPase/cinobufagin simulations highlight the same observations as described for the bufalin systems (Fig. 5B).

In short, in the MD simulations K^+^ was always surrounded by five complexing agents belonging to both αM1-4 and αM5-10 bundles. Complexing agents of a single Na^+^ included one residue from the αM1-4 bundle and two residues from the αM5-10 bundle, while Mg^2+^ exclusively coordinated to αM5-10 residues. Thus, K^+^ optimally spanned the separate bundles, Na^+^ also spanned the bundles but *via* fewer residues, while Mg^2+^ associated only with one helix bundle. Thus, the αM1-4/αM5-10 interface, and the contact surface for CTS binding, may vary depending on the ion. As the bufadienolides establish primarily hydrophobic interactions with the residues in the site, an increase in contact area is a main contributor for improving affinity. Therefore, we considered the calculation of the enzyme/bufadienolide surface overlap (i.e. contact area) as an indication of the indirect influence exerted by cations.

It appeared, that the overall contact surface for bufalin in Na^+^,K^+^-ATPase was increased in the presence of any cation compared to the system without ions. The calculations suggested that bufalin’s interactions with the binding site are weaker in the case of the least constrained site (Fig. S6), but the calculations were not sensitive enough to reveal differences between the ions. In an attempt to locate hotspots for the ion-induced changes in local conformations of the binding site, the extent of bufalin contacts with selected amino acids (based on Qiu et al. (2005)) as well as with individual helices (αM1-6) were calculated. Unfortunately, the detected fluctuations were not sufficiently systemdependent, possibly due to the limited time scale of the simulations. The importance of an ion for cross-linking the site remains hypothetical and awaits further investigation. However, certain observations from the MD simulations speak in favor of the ion-mediated confinement of the binding site. E.g. bufalin unbinding from the *apo* enzyme occurred *via* a semi-stable tilted binding mode in which the lactone was above the short αM6 helix and the α-surface was facing the extracellular solvent (Fig. S4B). This tilting was associated with an outward motion of αM1 and αM2 away from the αM5-10 bundle. With K^+^ in the ion sites, the helices would be restrained from such an outward movement and bufalin unbinding will be hindered.

The results of the MD simulations allow assignment of structural characteristics to the complexes with certain kinetic behaviors. It seems that the tilted binding mode represents the fastdissociating complex in biochemical experiments, and the coordination of both helix bundles by K^+^ is responsible for the disappearance of the fast-dissociating complex. K^+^ alone, but not Na^+^, Mg^2+^, or NMG^+^, was able to ensure homogeneity of the bufadienolide complexes in binding-dissociation studies, and Na^+^ and Mg^2+^ were not able to stabilize the helix bundles to the same extent as K^+^ in the MD simulations.

Thus, the *in silico* study of bufadienolide interactions with Na^+^,K^+^-ATPase performed in parallel with their biochemical characterization with Na^+^,K^+^-ATPase provided an explanation for the unexpected and specific effects of K^+^ in these reactions. As a specific ligand for the enzyme, K^+^ has high affinity for the extracellular cation transport sites accessible in the E2P form. The same conformation has high affinity for CTSs. While the presence of K^+^ decreases the affinity for cardenolide binding, it has a complex effect on bufadienolides. The crystal structure of the enzyme/bufalin complex (Laursen et al., 2015) revealed electrostatic interactions between K^+^ and the bufadienolide lactone ring. The K^+^-induced improvement in affinity, however, turned out to be valid for bufalin (and possibly for its derivatives) but not for all bufadienolides. The *in silico* calculations allowed us to look at two aspects of K^+^ presence in the extracellular sites: the direct effect on bufadienolide binding due to lactone/K^+^ coordination, and the indirect effect mediated by changes in protein structure. K^+^ turned out to be the only cation able to optimally span the two helix bundles and stabilize a well-defined local conformation of the CTS binding site. Therefore, K^+^ promotes formation of a homogeneous pool of bufadienolide complexes. For bufalin, this pre-formed site has an additional advantage: an optimal arrangement for electrostatic interactions between the lactone ring and K^+^, which improves affinity. Bufadienolides with more substituents on the steroid core (e.g. cinobufagin) do not gain from K^+^ presence in terms of affinity. On the contrary, steric clashes of the substituents hinder deep binding within the site and thus decrease affinity despite partial utilization of the electrostatic attraction between the ion and lactone carbonyl oxygen.

The structural fluctuations within the site are also reflected in the heterogeneity of Na^+^,K^+^-ATPase complexes with aglycones from the cardenolide family (Laursen et al., 2015). Cations occupying the extracellular sites seemed to influence the process as well (4). The cause of K^+^/cardenolide antagonism has been described on the basis of crystal structures (Laursen et al., 2013), and the cause of K^+^/bufadienolide agonism is described herein. The question of how glycosylation affects CTS binding is a subject for our on-going *in silico* study.

## Materials and Methods

### Biochemical characterization of E2Pi-bufadienolide complex of Na^+^,K^+^-ATPase

#### Enzyme preparation

Purified pig kidney Na^+^,K^+^-ATPase was prepared as previously described (Klodos et al., 2002). The specific ATPase activity of the preparation was about 1800 μmol Pi/hour per mg protein at 37 °C.

### Binding of bufadienolides to the E2Pi conformational state of Na^+^,K^+^-ATPase

The effect of cations on bufadienolides interactions with E2Pi state was estimated from the residual Na^+^,K^+^-ATPase activity after preincubation of bufadienolides with the enzyme essentially as described by Yatime et al. (2011) for ouabain interactions. The data were analyzed as in Yatime et al. (2011) using the commercial program KyPlot 5 (Kyence Inc.).

### Rate constants for bufalin dissociation from the E2Pi

Kinetics of bufalin dissociation from its complex with E2Pi was recorded on a SPEX Fluorolog-3 spectrofluorometer equipped with a thermostated cell compartment and a magnetic stirrer was used in a kinetic mode: excitation wavelength 370 nm (bandpass 5 nm), emission wavelength 485 nm (bandpass 5 nm). E2Pi-bufalin complex formed by incubation of the 33 μg/mL enzyme with 0.13 μM bufalin in 20 mM histidine pH 7.0, 4 mM H3PO4 (adjusted with N-methyl-D-glutamine), 4 mM MgCl2 in the absence or presence of 10 mM KCl at room temperature overnight. Then, the 100 μL aliquots were diluted in the 3 mL cuvettes with the same media but containing either 1.5 μM anthroylouabain or 1.5 μM anthroylouabain and 1 mM ouabain. The dissociation of bufalin is considered irreversible due to the presence of anthroylouabain (the concentration exceeds that of bufalin in the cuvette by a factor of 375). 1 mM ouabain prevents binding of both ligands and this sample thus serves as control for non-specific anthroylouabain binding and stability of its fluorescence in time.

### Computational protein and ligand preparation

The protein model was prepared from PDB entry 4RES (Laursen et al., 2015) (α1β1γ subunits from pig) using the Protein Preparation Wizard (Sastry et al., 2013) in Maestro (Schrödinger Suite 2019, Schrödinger LLC, New York, NY). This entailed capping of termini, addition of hydrogen atoms, potential flipping of His, Gln, and Asn, and a restrained minimization of the protein (max. 0.3 Å RMSD for heavy atoms). The protonation states of titratable residues were assessed by PROPKA 3 (Olsson et al., 2011), however, the protonation states of residues in the ion binding site were manually adjusted to be in accord with experiments by the Roux lab (Rui et al., 2016; Yu et al., 2011). The resulting model included disulfide bridges between CysB126-CysB149, CysB159-CysB175, and CysB213-CysB276; neutral AspA808, AspA926, GluA244, GluA327, GluA779, and GluA954; histidines HisB212, HisA286, HisA517, HisA550, HisA613, HisA659, HisA678, HisA875, and HisA912 were modeled as the ε-tautomer; and AspA369 was phosphorylated, while all other residues were modeled in the default state.

The chemical structure of bufalin was extracted from PDB entry 4RES (Laursen et al., 2015), while cinobufagin was extended manually from the bufalin structure using the build panel available in Maestro (Schrödinger Suite 2019). Both compounds were minimized using a conjugate gradient algorithm in 5000 steps and submitted to a conformational search by using a mixed torsional/low mode sampling algorithm as implemented in MacroModel (Schrödinger Suite 2019). The lowest energy conformation of each compound was used in the docking calculation and for force field parameter generation.

### Docking calculations

All docking calculations were performed using the IFD protocol (Sherman et al., 2006) employing Glide and Prime (Schrödinger Suite 2019). In the initial docking all vdW interactions were scaled to 50%, the SP level was applied, and a maximum of 200 poses were allowed. The centroid of the binding site was defined based on the co-crystallized bufalin (PDB ID: 4RES (Laursen et al., 2015)). In the optimization step, residues within 5 Å of the ligand were subjected to side chain optimization. The final docking step was performed in XP, and a maximum of 100 poses with associated energies within 30 kcal/mol of the lowest energy pose were reported in the results. The resulting poses were clustered based on their in-place conformation using the conformer cluster script available in Maestro (Schrödinger Suite 2019).

### System building for MD simulations

Each simulated system contains one Na^+^,K^+^-ATPase, one ligand, 1-2 structural cations, a POPC membrane patch, solvent and 0.2 M KCl. The system was built from scratch using a combined coarse grain/atomistic (CG/AA) approach as outlined below in the case of the Na^+^,K^+^-ATPase/bufalin/2xK^+^ system, while in the remaining systems the protein with bound ligand and structural ions were simply swapped into the first atomistic system before being equilibrated (Table 1).

**Table 1.** Overview of simulation protocol.

|  | Equilibration |  |  |  | Production run |
| --- | --- | --- | --- | --- | --- |
| Chronological step | 1 | 2 | 3 | 4 | 5 |
| Resolution | CG | CG | AA | AA | AA |
| Ensemble | NVT | NPT | NVT | NPT | NPT |
| Duration | 2 ns | 5 ns | 0.5 ns | 1 ns | 500 ns |
| Timestep | 10 fs | 10 fs | 2 fs | 2 fs | 2 fs |
| Position restraints | Protein/ligand/structural ions | Protein/ligand/structural ions | Protein/ligand/structural ions | Protein/ligand/structural ions | None |
| Thermostat | Berendsen | Velocity rescale | Nose-Hoover | Nose-Hoover | Nose-Hoover |
| Barostat | - | Berendsen | - | Parrinello-Rahman | Parrinello-Rahman |

The Na^+^,K^+^-ATPase/bufalin/2xK^+^ system was aligned onto the 4RES OPM (Lomize et al., 2012) structure to center the protein in coordinate space. A CG POPC membrane was then built around the protein in the xy plane using Insane (Wassenaar et al., 2015) and Martinize tools available from the Marrink group’s homepage^1^. The system was solvated and neutralized with 0.2 M KCl before being minimized and equilibrated according to step 1 and 2 in Table 1. The system was then converted into its atomistic equivalent using the Backward tool (Wassenaar et al., 2014). In order to ensure that the starting point of the AA simulations was exactly as intended during protein preparation, the backmapped protein was replaced by the original Na^+^,K^+^-ATPase/ligand/ion complex. The atomistic system was then minimized using a conjugate gradient algorithm and further equilibrated in AA resolution before production runs, as outlined in step 3 to 5 in Table 1. As we are interested in the ligand dynamics and stability within the protein we perform data analysis immediately following release of position restraints. Each system repeat was equilibrated separately to ensure maximal sampling of each molecular system. In all simulations, the RMSD of Cα atoms was observed to converge almost immediately (Fig. S7)

### Molecular dynamics simulations

All simulations were performed in Gromacs 2019.2 using periodic boundary conditions. For the CG simulations, the MARTINI 2.2 force field was used (de Jong et al., 2013; Marrink et al., 2007; Monticelli et al., 2008). The vdW interactions were treated using cut-offs at 11 Å and the potential-shift-Verlet modifier, while electrostatic interactions were treated using the reaction-field method cutoff at 11 Å and a dielectric constant of 0 (= infinite) beyond the cut-off. The neighbor list was maintained using Verlet buffer lists. Temperature was kept at 310 K and pressure at 1 bar. The AA resolution simulations were performed using the CHARMM36m force field (Huang et al., 2017; Klauda et al., 2010; MacKerell et al., 1998), the TIPS3P water model (Durell et al., 1994), and CHARMM-compatible ligand parameters as described below. The vdW interactions were treated by cut-offs at 12 Å and a force-switch modifier after 10 Å, while electrostatic interactions were treated using PME. The neighbor list was maintained using Verlet buffer lists, and bonds linking hydrogen atoms to heavy atoms were restrained using LINCS (Hess, 2008). The temperature was maintained at 310 K using a coupling constant of 1, and pressure was maintained at 1 bar using a coupling constant of 4, a compressibility factor of 4.5 x10^-5^ and semiisotropic coupling to xy and z dimensions separately.

### Force field parameters for ligands and phosphorylated aspartate

Force field parameters for bufalin and cinobufagin were obtained by analogy using the ParamChem webserver (Vanommeslaeghe et al., 2012) and the CHARMM generalized force field (Vanommeslaeghe et al., 2010). The associated penalties were assessed and indicated the parameters were a good fit. The applied parameters can be found in supporting data file S1-S4. The CHARMM compatible parameters for the phosphorylated aspartate were taken from Damjanovic et al. (2009).

### Calculation of enzyme/bufadienolide contacts

The interatomic contact surface between bufadienolides and Na^+^,K^+^-ATPase was calculated using the dr_sasa software (Ribeiro et al., 2019) in mode 4. Mode 4 calculates the contact surface (i.e. overlapping surface area between ligand and protein) based on close atom-atom interactions and thus not by using a spherical water probe as done commonly. This ensures a more thorough calculation of the intramolecular contacts. The calculation was performed for every 100th frame of each trajectory i.e. a frame per every 40 ps.

### Computational analyses

All analyses other than the contact calculations were performed using in-house scripts. All figures of molecular systems were made using VMD 1.9.3 or ChemDraw.

## Acknowledgments

This work was supported by grants from the Danish Council for Independent Research (DFF-7016-00125, to N.U.F.), the A.P. Møller Foundation for the Advancement of Medical Science (to

N.U.F.), and Novo Nordic Foundation (NNF18OC0032608, to L.K.L.). We are thankful to B. Bjerring Jensen for excellent technical assistance. All computations were performed at the Grendel cluster (http://www.cscaa.dk/grendel/hardware/) through the Centre for Scientific Computing - Aarhus.

## Author Contributions

All authors participated in the research design and data analyses; L.K.L performed the *in silico* experiments; N.U.F. performed the *in vitro* experiments. All authors contributed to the writing of the paper.

## Footnotes

1 http://cgmartini.nl/index.php/tools2/proteins-and-bilayers

## References

Damjanovic, A., Garcia-Moreno, B., & Brooks, B. R. (2009). Self-guided Langevin dynamics study of regulatory interactions in NtrC. Proteins-Structure Function and Bioinformatics, 76(4), 1007–1019. doi:10.1002/prot.22439

de Jong, D. H., Singh, G., Bennett, W. F., Arnarez, C., Wassenaar, T. A., Schafer, L. V., Periole, X., Tieleman, D. P., & Marrink, S. J. (2013). Improved parameters for the MARTINI coarse-grained protein force field. Journal of Chemical Theory and Computation, 9(1), 687–697. doi:10.1021/ct300646g

Durell, S. R., Brooks, B. R., & Bennaim, A. (1994). Solvent-induced forces between 2 hydrophilic groups. Journal of Physical Chemistry, 98(8), 2198–2202. doi:DOI 10.1021/j100059a038

Fricke, U., & Klaus, W. (1981). The influence of reduced serum potassium level on the toxicity of some cardenolides in guinea-pigs. Basic Research in Cardiology, 76(1), 62–78. doi:Doi 10.1007/Bf01908163

Gelbart, A., Hall, R. J., & Goldman, R. (1980). Effects of hypokalemia on the cardiotropic actions of digoxin in dogs - correlation with inhibition of cardiac Na+,K+-adenosine triphosphatase. Circulation Research, 46(6), I173–I174.

Gregersen, J. L., Mattle, D., Fedosova, N. U., Nissen, P., & Reinhard, L. (2016). Isolation, crystallization and crystal structure determination of bovine kidney Na+, K+-ATPase. Acta Crystallographica Section F-Structural Biology Communications, 72, 282–287. doi: 10.1107/S2053230x1600279x

Hansen, O., & Skou, J. C. (1973). A study on the influence of the concentration of Mg2+, Pi, K+, Na+, and Tris on (Mg2+ + Pi)-supported g-strophanthin binding to (Na+ = K+) activated ATPase from ox brain. Biochimica et Biophysica Acta (BBA) - Bioenergetics, 311(1), 51–66. doi:10.1016/0005-2736(73)90254-x

Hess, B. (2008). P-LINCS: a parallel linear constraint solver for molecular simulation. Journal of Chemical Theory and Computation, 4(1), 116–122. doi:10.1021/ct700200b

Huang, J., Rauscher, S., Nawrocki, G., Ran, T., Feig, M., de Groot, B. L., Grubmuller, H., & MacKerell, A. D. (2017). CHARMM36m: an improved force field for folded and intrinsically disordered proteins. Nature Methods, 14(1), 71–73. doi:10.1038/Nmeth.4067

Katz, A., Lifshitz, Y., Bab-Dinitz, E., Kapri-Pardes, E., Goldshleger, R., Tal, D. M., & Karlish, S. J. D. (2010). Selectivity of Digitalis Glycosides for Isoforms of Human Na,K-ATPase. Journal of Biological Chemistry, 285(25), 19582–19592. doi:10.1074/jbc.M110.119248

Katz, A., Tal, D. M., Heller, D., Habeck, M., Ben Zeev, E., Rabah, B., Bar Kana, Y., Marcovich, A. L., & Karlish, S. J. D. (2015). Digoxin derivatives with selectivity for the alpha 2 beta 3 isoform of Na,K-ATPase potently reduce intraocular pressure. Proceedings of the National Academy of Sciences of the United States of America, 112(44),13723–13728. doi:10.1073/pnas.1514569112

Klauda, J. B., Venable, R. M., Freites, J. A., O’Connor, J. W., Tobias, D. J., Mondragon-Ramirez, C., Vorobyov, I., MacKerell, A. D., Jr., & Pastor, R. W. (2010). Update of the CHARMM allatom additive force field for lipids: validation on six lipid types. Journal of Physical Chemistry B, 114(23), 7830–7843. doi:10.1021/jp101759q

Klodos, I., Esmann, M., & Post, R. L. (2002). Large-scale preparation of sodium-potassium ATPase from kidney outer medulla. Kidney International, 62(6), 2097–2100. doi:DOI 10.1046/j.1523-1755.2002.00654.x

Laursen, M., Gregersen, J. L., Yatime, L., Nissen, P., & Fedosova, N. U. (2015). Structures and characterization of digoxin- and bufalin-bound Na+, K+-ATPase compared with the ouabain-bound complex. Proceedings of the National Academy of Sciences of the United States of America, 112(6), 1755–1760. doi:10.1073/pnas.1422997112

Laursen, M., Yatime, L., Nissen, P., & Fedosova, N. U. (2013). Crystal structure of the high-affinity Na+,K+-ATPase-ouabain complex with Mg2+ bound in the cation binding site. Proceedings of the National Academy of Sciences of the United States of America, 110(27), 10958–10963. doi:10.1073/pnas.1222308110

Lomize, M. A., Pogozheva, I. D., Joo, H., Mosberg, H. I., & Lomize, A. L. (2012). OPM database and PPM web server: resources for positioning of proteins in membranes. Nucleic Acids Research, 40(D1), D370–D376. doi:10.1093/nar/gkr703

MacKerell, A. D., Bashford, D., Bellott, M., Dunbrack, R. L., Evanseck, J. D., Field, M. J., Fischer, S., Gao, J., Guo, H., Ha, S., Joseph-McCarthy, D., Kuchnir, L., Kuczera, K., Lau, F. T. K., Mattos, C., Michnick, S., Ngo, T., Nguyen, D. T., Prodhom, B., Reiher, W. E., Roux, B., Schlenkrich, M., Smith, J. C., Stote, R., Straub, J., Watanabe, M., Wiorkiewicz-Kuczera, J., Yin, D., & Karplus, M. (1998). All-atom empirical potential for molecular modeling and dynamics studies of proteins. Journal of Physical Chemistry B, 102(18), 3586–3616. doi:DOI 10.1021/jp973084f

Marrink, S. J., Risselada, H. J., Yefimov, S., Tieleman, D. P., & de Vries, A. H. (2007). The MARTINI force field: coarse grained model for biomolecular simulations. Journal of Physical Chemistry B, 111(27), 7812–7824. doi:10.1021/jp071097f

Monticelli, L., Kandasamy, S. K., Periole, X., Larson, R. G., Tieleman, D. P., & Marrink, S. J. (2008). The MARTINI coarse-grained force field: extension to proteins. Journal of Chemical Theory and Computation, 4(5), 819–834. doi:10.1021/ct700324x

Ogawa, H., Shinoda, T., Cornelius, F., & Toyoshima, C. (2009). Crystal structure of the sodium-potassium pump (Na+,K+-ATPase) with bound potassium and ouabain. Proceedings of the National Academy of Sciences of the United States of America, 106(33), 13742–13747. doi:10.1073/pnas.0907054106

Olsson, M. H. M., Søndergaard, C. R., Rostkowski, M., & Jensen, J. H. (2011). PROPKA3: consistent treatment of internal and surface residues in empirical pKa predictions. Journal of Chemical Theory and Computation, 7(2), 525–537. doi:10.1021/ct100578z

Qiu, L. Y., Krieger, E., Schaftenaar, G., Swarts, H. G., Willems, P. H., De Pont, J. J., & Koenderink, J. B. (2005). Reconstruction of the complete ouabain-binding pocket of Na,K-ATPase in gastric H,K-ATPase by substitution of only seven amino acids. Journal of Biological Chemistry, 280(37), 32349–32355. doi:10.1074/jbc.M505168200

Ribeiro, J., Rios-Vera, C., Melo, F., & Schuller, A. (2019). Calculation of accurate interatomic contact surface areas for the quantitative analysis of non-bonded molecular interactions. Bioinformatics, 35(18), 3499–3501. doi:10.1093/bioinformatics/btz062

Rui, H., Artigas, P., & Roux, B. (2016). The selectivity of the Na+/K+-pump is controlled by binding site protonation and self-correcting occlusion. Elife,5. doi:ARTN e16616 10.7554/eLife.16616

Sandtner, W., Egwolf, B., Khalili-Araghi, F., Sanchez-Rodriguez, J. E., Roux, B., Bezanilla, F., & Holmgren, M. (2011). Ouabain binding site in a functioning Na+/K+ ATPase. Journal of Biological Chemistry, 286(44), 38177–38183. doi:10.1074/jbc.M111.267682

Sastry, G. M., Adzhigirey, M., Day, T., Annabhimoju, R., & Sherman, W. (2013). Protein and ligand preparation: parameters, protocols, and influence on virtual screening enrichments. Journal of Computer-Aided Molecular Design, 27(3), 221–234. doi:10.1007/s10822-013-9644-8

Sherman, W., Day, T., Jacobson, M. P., Friesner, R. A., & Farid, R. (2006). Novel procedure for modeling ligand/receptor induced fit effects. Journal of Medicinal Chemistry, 49(2), 534–553. doi:10.1021/jm050540c

Vanommeslaeghe, K., Hatcher, E., Acharya, C., Kundu, S., Zhong, S., Shim, J., Darian, E., Guvench, O., Lopes, P., Vorobyov, I., & Mackerell, A. D., Jr. (2010). CHARMM general force field: a force field for drug-like molecules compatible with the CHARMM all-atom additive biological force fields. Journal of Computational Chemistry, 31(4), 671–690. doi:10.1002/jcc.21367

Vanommeslaeghe, K., Raman, E. P., & MacKerell, A. D. (2012). Automation of the CHARMM General Force Field (CGenFF) II: Assignment of Bonded Parameters and Partial Atomic Charges. Journal of Chemical Information and Modeling, 52(12),3155–3168. doi:10.1021/ci3003649

Varma, S., & Rempe, S. B. (2006). Coordination numbers of alkali metal ions in aqueous solutions. Biophysical Chemistry, 124(3), 192–199. doi:10.1016/j.bpc.2006.07.002

Wassenaar, T. A., Ingolfsson, H. I., Bockmann, R. A., Tieleman, D. P., & Marrink, S. J. (2015). Computational lipidomics with insane: a versatile tool for generating custom membranes for molecular simulations. Journal of Chemical Theory and Computation, 11(5), 2144–2155. doi:10.1021/acs.jctc.5b00209

Wassenaar, T. A., Pluhackova, K., Bockmann, R. A., Marrink, S. J., & Tieleman, D. P. (2014). Going backward: a flexible geometric approach to reverse transformation from coarse grained to atomistic models. Journal of Chemical Theory and Computation, 10(2), 676–690. doi:10.1021/ct400617g

Weigand, K. M., Laursen, M., Swarts, H. G. P., Engwerda, A. H. J., Prufert, C., Sandrock, J., Nissen,P., Fedosova, N. U., Russel, F. G. M., & Koenderink, J. B. (2014). Na+,K+-ATPase isoform selectivity for digitalis-like compounds is determined by two amino acids in the first extracellular loop. Chemical Research in Toxicology, 27(12), 2082–2092. doi:10.1021/tx500290k

Yatime, L., Laursen, M., Morth, J. P., Esmann, M., Nissen, P., & Fedosova, N. U. (2011). Structural insights into the high affinity binding of cardiotonic steroids to the Na+,K+-ATPase. Journal of Structural Biology, 174(2), 296–306. doi:10.1016/j.jsb.2010.12.004

Yu, H. B., Ratheal, I. M., Artigas, P., & Roux, B. (2011). Protonation of key acidic residues is critical for the K+-selectivity of the Na/K pump. Nature Structural & Molecular Biology, 18(10), 1159–U1116. doi:10.1038/nsmb.2113

Yu, H. B., Whitfield, T. W., Harder, E., Lamoureux, G., Vorobyov, I., Anisimov, V. M., MacKerell, A. D., & Roux, B. (2010). Simulating monovalent and divalent ions in aqueous solution using a drude polarizable force field. Journal of Chemical Theory and Computation, 6(3), 774–786. doi:10.1021/ct900576a

Zheng, H., Chruszcz, M., Lasota, P., Lebioda, L., & Minor, W. (2008). Data mining of metal ion environments present in protein structures. Journal of Inorganic Biochemistry, 102(9), 1765–1776. doi:10.1016/j.jinorgbio.2008.05.006

