## Supplementary Material for "Beneficent and maleficent effects of cations on bufadienolide binding to Na^+^,K^+^-ATPase"

Contents:

Supporting table S1. Effect of cations on bufalin and cinobufagin binding.
Supporting table S2. Induced fit docking scores.
Supporting figure S1. Cation effect on bufadienolide interactions with $Na^+,K^+$-ATPase as reflected by enzyme inhibition.
Supporting figure S2. Major bufalin binding clusters from docking calculation.
Supporting figure S3. Major cinobufagin binding clusters from docking calculation.
Supporting figure S4. Bufalin unbinding.
Supporting figure S5. RMSD of the ion in site II.
Supporting figure S6. Enzyme/bufadienolide contacts.
Supporting figure S7. Enzyme RMSD.
Supporting data file S1. Bufalin itp file.
Supporting data file S2. Bufalin prm file.
Supporting data file S3. Cinobufagin itp file.
Supporting data file S4. Cinobufagin prm file.

**Supporting table S1.** Effect of cations on bufalin and cinobufagin binding. The basic media always contains 3 mM $MgCl_2$.

| | Bufalin | | | | | | | | | Cinobufagin | |
|---|---|---|---|---|---|---|---|---|---|---|---|
| | No ions (3 mM $Mg^{2+}$ ) | $Mg^{2+}$ | $Na^{+}$ | | $K^{+}$ | | | | $NMG^{+}$ | No ions (3 mM $Mg^{2+}$ ) | $K^{+}$ |
| | | 9 mM | 100 mM | 200 mM | 1 mM | 10 mM | 100 mM | 200 mM | 200 mM | | 200 mM |
| $K_d$ (nM) | 12 | 11 | 80 | 600 | 16 | 8 | 5 | 11 | 11 | 51 | 103 |
| ±SE | 6 | 0.9 | 13 | 88 | 7 | 4 | 4 | 4 | 5 | 19 | 12 |
| Stable complex (%) | 83 | 83 | 84 | 87 | 95 | 100 | 100 | 100 | 84 | 88 | 100 |

**Supporting table S2.** Resulting scores of the IFD calculations of bufalin and cinobufagin into $Na^+$,$K^+$-ATPase with and without $K^+$ ions. All scores are reported as averages in kcal/mol with the associated standard deviations (SD).

| Cluster | n | Av. XP Gscore | SD | Av. IFDScore | SD |
|---|---|---|---|---|---|
| Bufalin with $K^+$ | | | | | |
| $B1_{K+}$ | 1 | -8.1 | - | -2729.6 | - |
| $B2_{K+}$ | 1 | -4.3 | - | -2725.6 | - |
| $B3_{K+}$ | 1 | -7.0 | - | -2728.2 | - |
| $B4_{K+}$ | 5 | -8.8 | 0.5 | -2729.9 | 0.4 |
| $B5_{K+}$ | 8 | -5.6 | 0.8 | -2726.8 | 0.8 |
| $B6_{K+}$ | 24 | -7.0 | 0.7 | -2728.6 | 0.9 |
| Bufalin without $K^+$ | | | | | |
| B1 | 1 | -7.4 | - | -2726.6 | - |
| B2 | 1 | -4.9 | - | -2724.6 | - |
| B3 | 1 | -5.4 | - | -2724.7 | - |
| B4 | 10 | -7.1 | 2.1 | -2726.5 | 2.1 |
| B5 | 10 | -6.2 | 0.8 | -2725.7 | 0.8 |
| B6 | 15 | -7.7 | 0.8 | -2727.6 | 1.0 |
| Cinobufagin with $K^+$ | | | | | |
| $C1_{K+}$ | 1 | -7.6 | - | -2730.9 | - |
| $C2_{K+}$ | 1 | -5.4 | - | -2729.1 | - |
| $C3_{K+}$ | 21 | -6.7 | 0.8 | -2730.5 | 1.0 |
| Cinobufagin without $K^+$ | | | | | |
| C1 | 20 | -6.3 | 0.9 | -2727.7 | 1.0 |
| C2 | 1 | -6.4 | - | -2727.5 | - |
| C3 | 14 | -5.7 | 0.7 | -2726.9 | 0.8 |
| C4 | 9 | -6.1 | 1.7 | -2727.5 | 1.6 |
| C5 | 8 | -6.7 | 1.4 | -2728.0 | 1.5 |
| C6 | 18 | -6.3 | 1.3 | -2727.6 | 1.4 |
| C7 | 1 | -5.2 | - | -2726.5 | - |

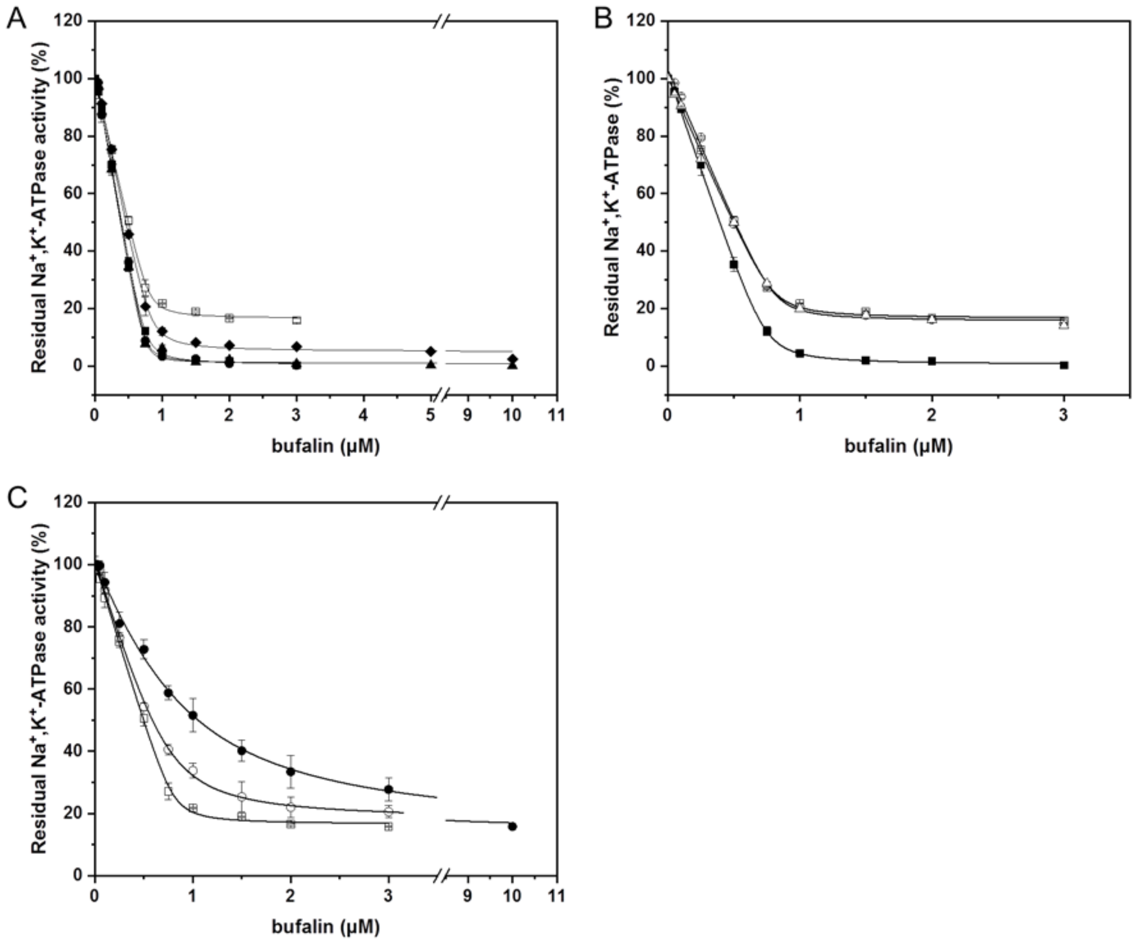


**Supporting Figure S1**. Cation effect on bufadienolide interactions with $Na^+$,$K^+$-ATPase as reflected by enzyme inhibition. **(A)** Bufalin binding in the presence of 0 mM KCl (□), 1mM KCl (⧫), 10 mM KCl (▲), 100 mM KCl (●), and 200 mM KCl (■). **(B)** Bufalin binding in the presence of 3 mM $MgCl_2$ (□), 3+9 mM $MgCl_2$ (Δ) or 200 mM NMGCl (○), and 200 mM KCl (■). **(C)** Bufalin binding in the presence of 0 mM NaCl (□), 100 mM NaCl (○), and 200 mM NaCl (●). All experiments were performed in triplicates in the presence of 3 mM $MgCl_2$.

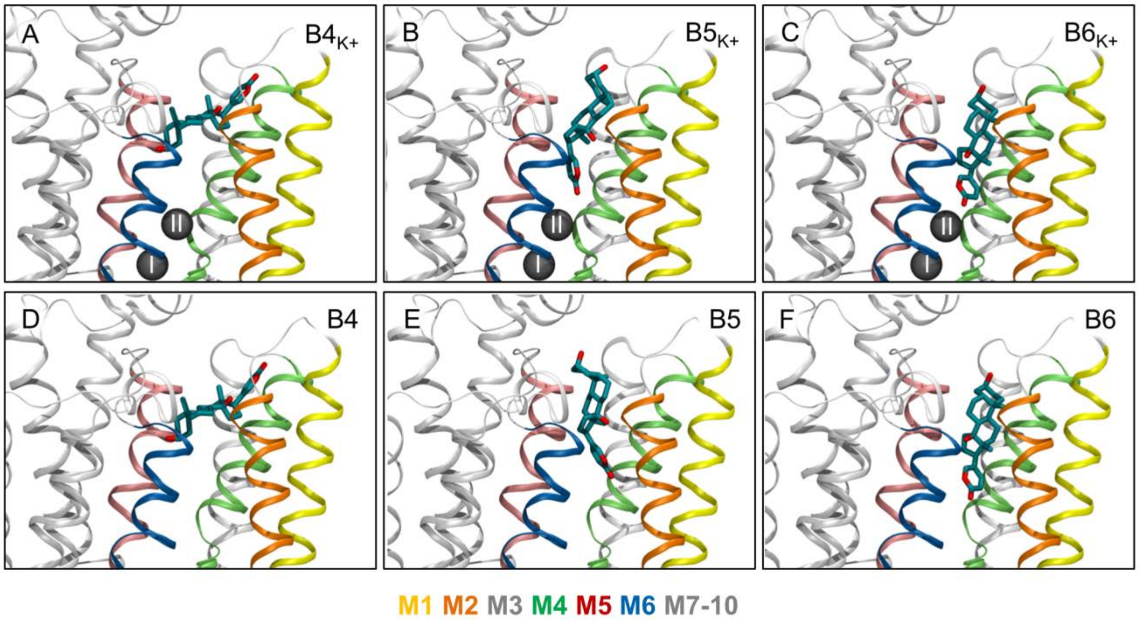


**Supporting figure S2**. Main binding modes of bufalin in $Na^+$,$K^+$-ATPase in the presence **(ABC)** or absence **(DEF)** of $K^+$ ions. Bufalin is shown in cyan, $K^+$ ions as gray spheres, and the enzyme as ribbons.

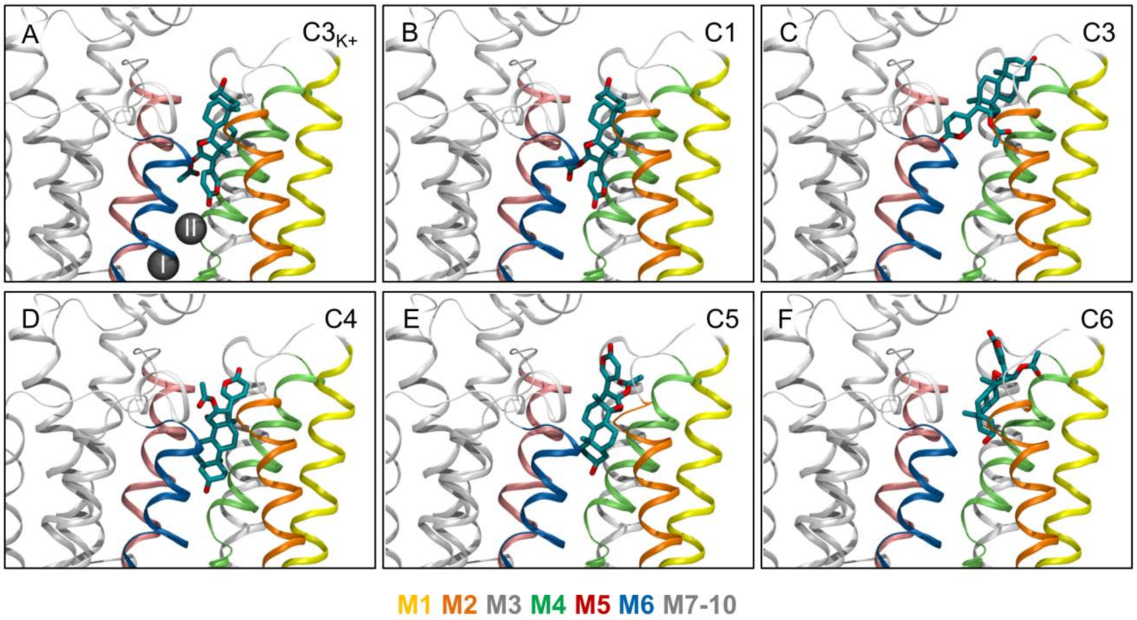


**Supporting figure S3**. Main binding modes of cinobufagin in $Na^+$,$K^+$-ATPase in the presence **(A)** or absence **(BCDEF)** of $K^+$ ions. Bufalin is shown in cyan, $K^+$ ions as gray spheres, and the enzyme as ribbons.

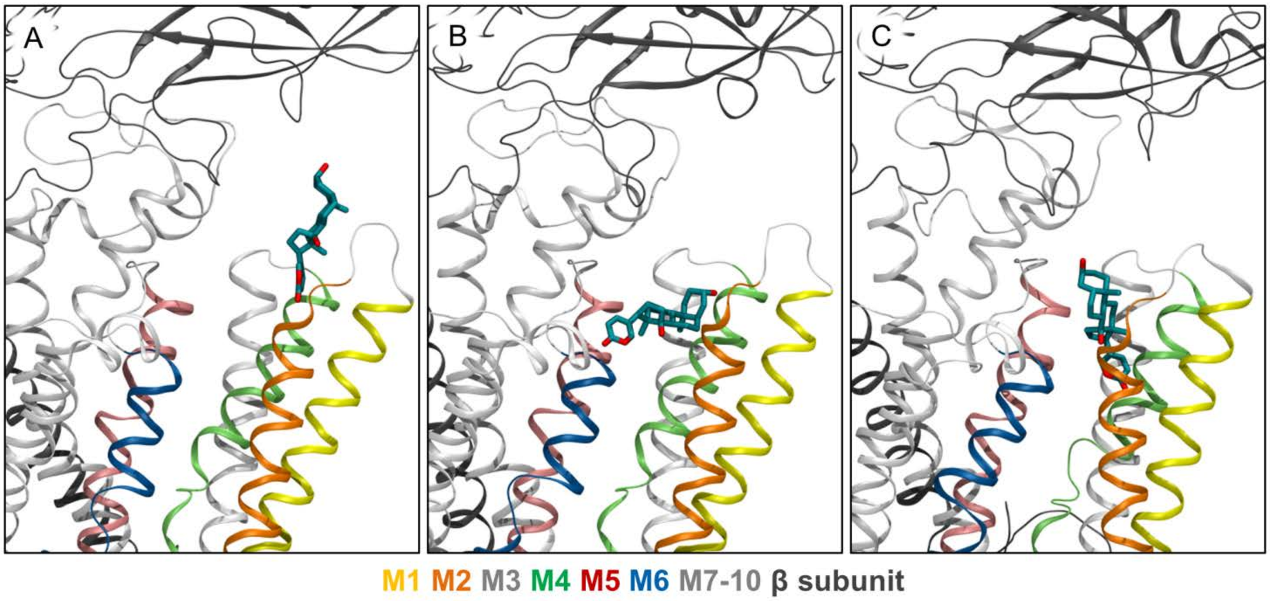


**Supporting figure S4**. Bufalin unbinding from apo $Na^+$,$K^+$-ATPase. **(A)** Snapshot of the most extracellular position of bufalin from MD1. **(B)** Snapshot of the tilted binding mode observed prior to further unbinding from $Na^+$,$K^+$-ATPase in MD1. This tilted binding mode was also observed in $C4_{buf,K+}$ and $C4_{buf}$ from the docking calculations. **(C)** Snapshot of the most extracellular position of bufalin from MD2. Bufalin is shown in cyan, $K^+$ ions as gray spheres, and the enzyme as ribbons.

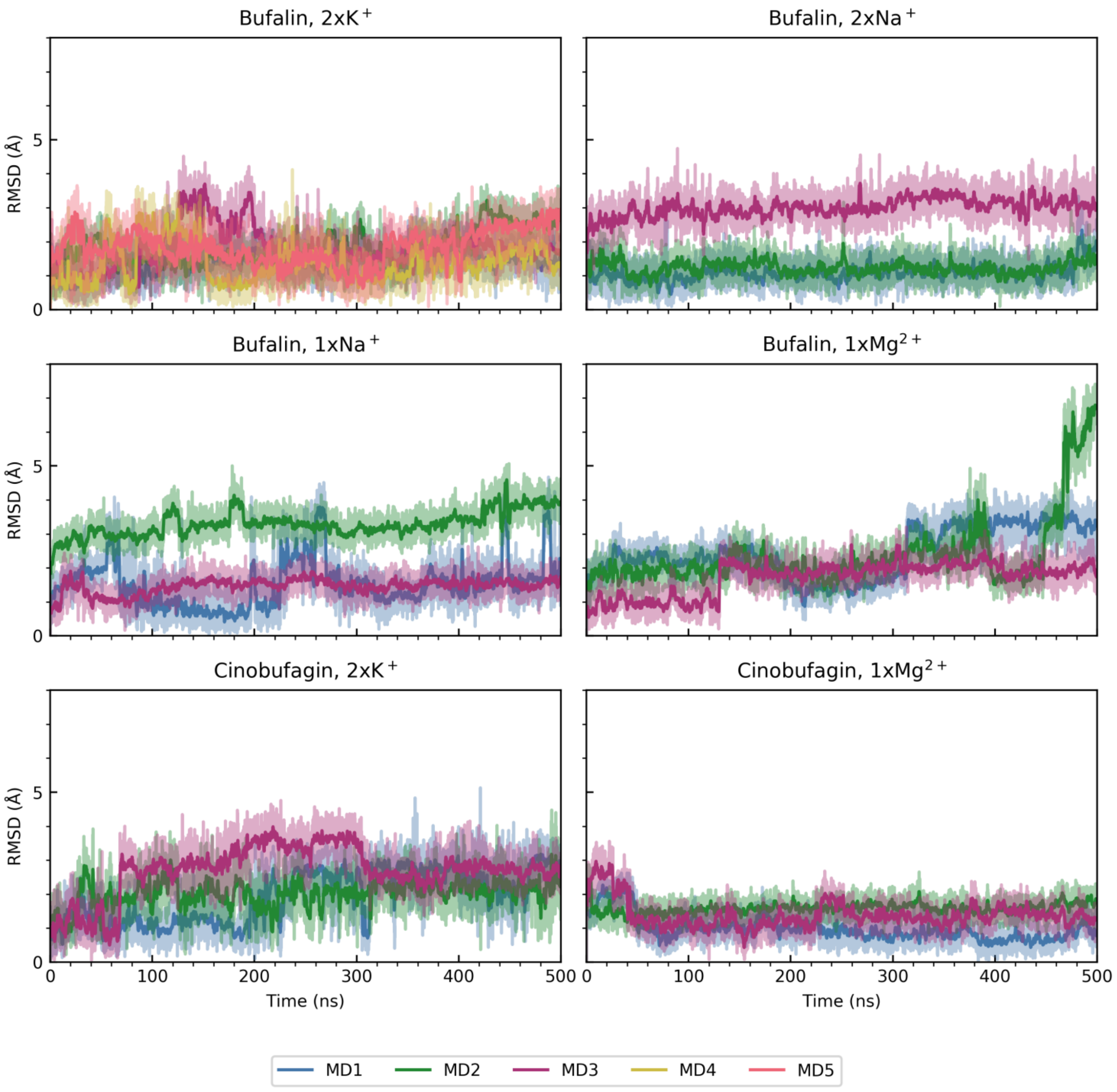


**Supporting figure S5**. The time progressed RMSD of the ion located in cation transport site II within the enzyme. All trajectories were aligned prior to analysis using M5-10 of the enzyme. The running average is shown in opaque hues while the raw data is shown in transparent hues based on the color legend below the plot.

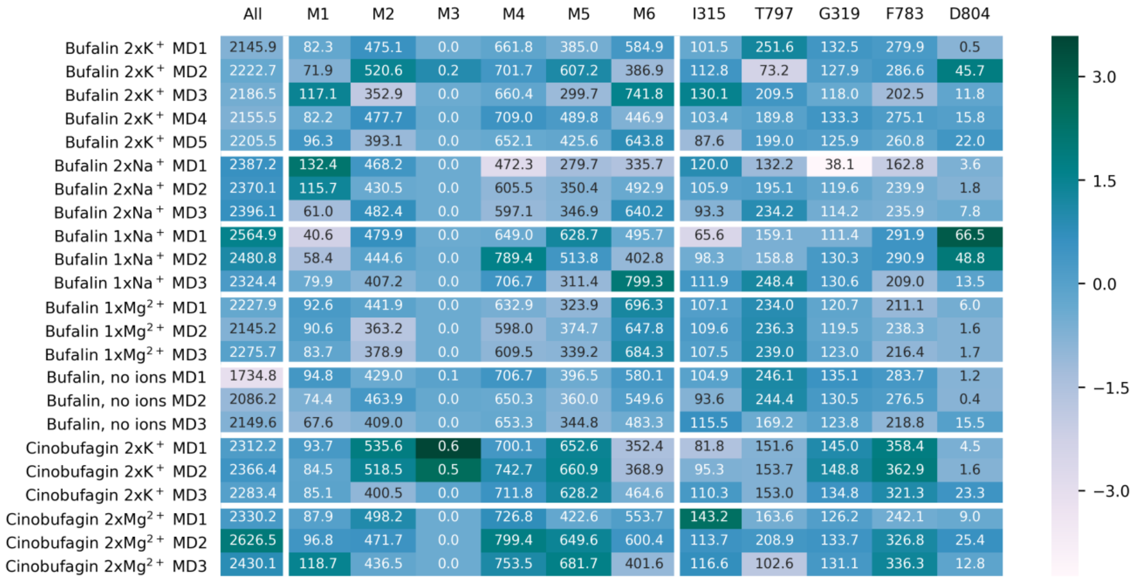


**Supporting figure S6.** Enzyme/bufadienolide contacts. Each row represents a simulation repeat while each column denotes the part of the enzyme included in the contact calculation. Each cell holds the average contact value in Å$^2$ for a given simulation repeat. The color of the cell relates the simulation repeat average to the overall average obtained across all molecular systems and repeats for that particular part of the enzyme. Thus, zero (light blue) on the color scale bar represents the average contact value of a given column, while green and cream represents contact values more than 3 times the standard deviation above and below the average, respectively. These extreme colors thus depict formation or disappearance of contacts compared to the average across all molecular systems i.e. a given column.

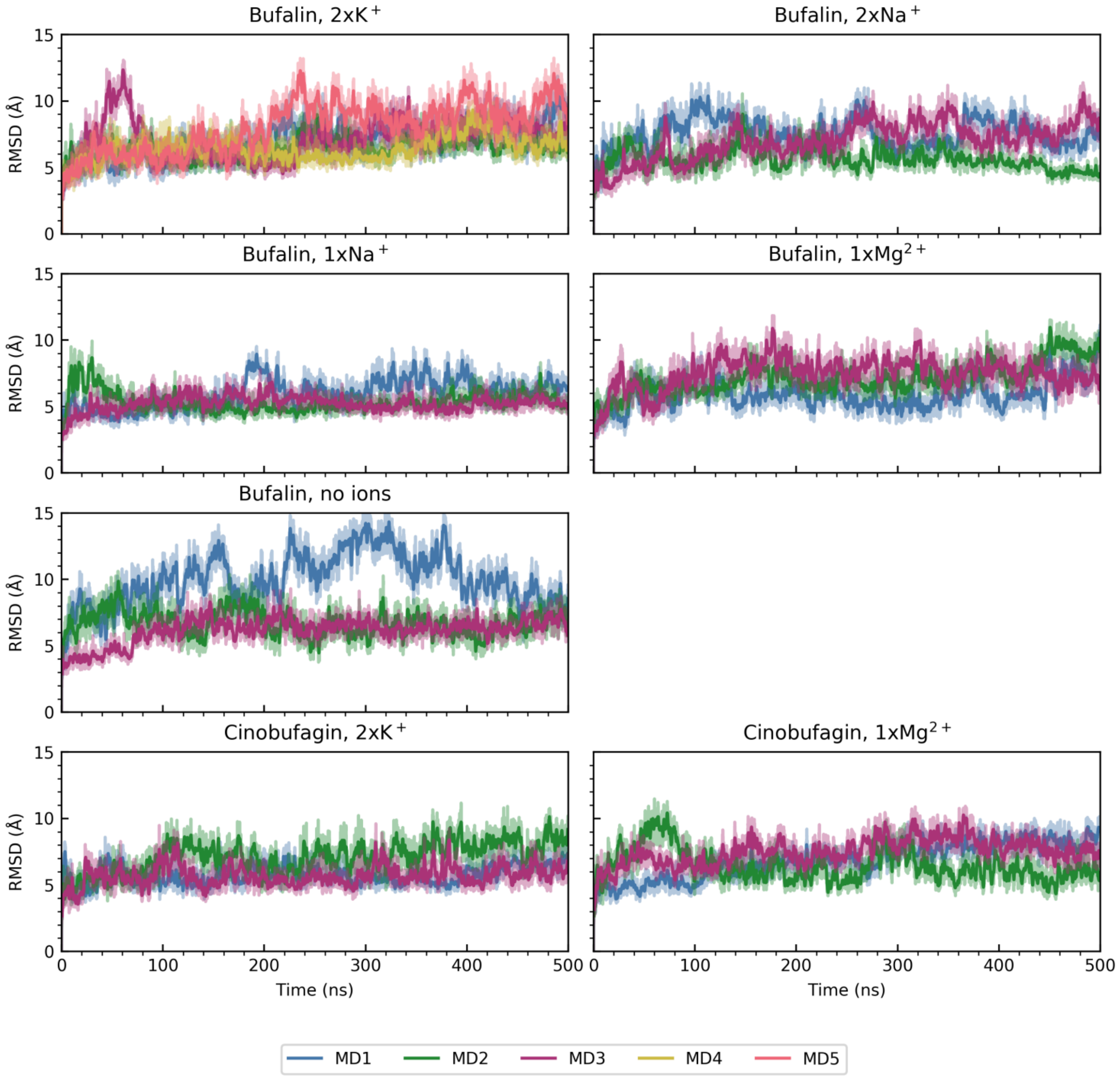


**Supporting figure S7**. The time progressed RMSD of all Cα atoms of the enzyme. All trajectories were aligned prior to analysis using M5-10 of the enzyme. The running average is shown in opaque hues while the raw data is shown in transparent hues based on the color legend below the plot.

**Supporting data file S1.** Bufalin itp file.

```
; Created by cgenff_charmm2gmx.py
 [ moleculetype ]
; Name  nrexcl
bufalin  3

[ atoms ]
; nr  type resnr residue  atom   cgnr   charge mass  typeB    chargeB massB
; residue 1 bufalin_axial  rtp bufalin_axial  q       qsum
     1      CG331      1 bufalin_axial     C18      1     -0.268     12.011   ;
     2     CG3RC1      1 bufalin_axial     C13      2      0.001     12.011   ;
     3      CG321      1 bufalin_axial     C12      3     -0.183     12.011   ;
     4      CG321      1 bufalin_axial     C11      4     -0.182     12.011   ;
     5      CG311      1 bufalin_axial      C9      5     -0.090     12.011   ;
     6      CG301      1 bufalin_axial     C10      6     -0.001     12.011   ;
     7      CG331      1 bufalin_axial     C19      7     -0.274     12.011   ;
     8      CG321      1 bufalin_axial      C1      8     -0.180     12.011   ;
     9      CG321      1 bufalin_axial      C2      9     -0.179     12.011   ;
    10      CG311      1 bufalin_axial      C3     10      0.135     12.011   ;
    11      OG311      1 bufalin_axial      O3     11     -0.650     15.999   ;
    12      CG321      1 bufalin_axial      C4     12     -0.177     12.011   ;
    13      CG311      1 bufalin_axial      C5     13     -0.090     12.011   ;
    14      CG321      1 bufalin_axial      C6     14     -0.181     12.011   ;
    15      CG321      1 bufalin_axial      C7     15     -0.181     12.011   ;
    16      CG311      1 bufalin_axial      C8     16     -0.090     12.011   ;
    17     CG3RC1      1 bufalin_axial     C14     17      0.238     12.011   ;
    18      OG311      1 bufalin_axial     O14     18     -0.651     15.999   ;
    19     CG3C52      1 bufalin_axial     C15     19     -0.181     12.011   ;
    20     CG3C52      1 bufalin_axial     C16     20     -0.177     12.011   ;
    21     CG3C51      1 bufalin_axial     C17     21     -0.039     12.011   ;
    22     CG2R62      1 bufalin_axial     C20     22      0.032     12.011   ;
    23     CG2R62      1 bufalin_axial     C22     23     -0.272     12.011   ;
    24     CG2R62      1 bufalin_axial     C23     24     -0.347     12.011   ;
    25     CG2R63      1 bufalin_axial     C24     25      0.490     12.011   ;
    26      OG2D4      1 bufalin_axial     O24     26     -0.460     15.999   ;
    27     OG3R60      1 bufalin_axial     O21     27     -0.362     15.999   ;
    28     CG2R62      1 bufalin_axial     C21     28      0.119     12.011   ;
    29       HGA3      1 bufalin_axial    H181     29      0.090      1.008   ;
    30       HGA3      1 bufalin_axial    H182     30      0.090      1.008   ;
    31       HGA3      1 bufalin_axial    H183     31      0.090      1.008   ;
    32       HGA2      1 bufalin_axial    H121     32      0.090      1.008   ;
    33       HGA2      1 bufalin_axial    H122     33      0.090      1.008   ;
    34       HGA2      1 bufalin_axial    H111     34      0.090      1.008   ;
    35       HGA2      1 bufalin_axial    H112     35      0.090      1.008   ;
    36       HGA1      1 bufalin_axial      H9     36      0.090      1.008   ;
    37       HGA3      1 bufalin_axial    H191     37      0.090      1.008   ;
```

```
   38      HGA3      1 bufalin_axial    H192     38      0.090      1.008   ;
   39      HGA3      1 bufalin_axial    H193     39      0.090      1.008   ;
   40      HGA2      1 bufalin_axial    H11     40      0.090      1.008   ;
   41      HGA2      1 bufalin_axial    H12     41      0.090      1.008   ;
   42      HGA2      1 bufalin_axial    H21     42      0.090      1.008   ;
   43      HGA2      1 bufalin_axial    H22     43      0.090      1.008   ;
   44      HGA1      1 bufalin_axial     H3     44      0.090      1.008   ;
   45      HGP1     1 bufalin_axial    HO3     45      0.419      1.008   ;
   46      HGA2      1 bufalin_axial    H41     46      0.090      1.008   ;
   47      HGA2      1 bufalin_axial    H42     47      0.090      1.008   ;
   48      HGA1      1 bufalin_axial     H5     48      0.090      1.008   ;
   49      HGA2      1 bufalin_axial    H61     49      0.090      1.008   ;
   50      HGA2      1 bufalin_axial    H62     50      0.090      1.008   ;
   51      HGA2      1 bufalin_axial    H71     51      0.090      1.008   ;
   52      HGA2      1 bufalin_axial    H72     52      0.090      1.008   ;
   53      HGA1      1 bufalin_axial     H8     53      0.090      1.008   ;
   54      HGP1     1 bufalin_axial    HO14     54      0.420      1.008   ;
   55      HGA2      1 bufalin_axial    H151     55      0.090      1.008   ;
   56      HGA2      1 bufalin_axial    H152     56      0.090      1.008   ;
   57      HGA2      1 bufalin_axial    H161     57      0.090      1.008   ;
   58      HGA2      1 bufalin_axial    H162     58      0.090      1.008   ;
   59      HGA1      1 bufalin_axial    H17     59      0.090      1.008   ;
   60     HGR62      1 bufalin_axial     H2     60      0.268      1.008   ;
   61     HGR62      1 bufalin_axial    H23     61      0.199      1.008   ;
   62     HGR62      1 bufalin_axial     H1     62      0.284      1.008   ;

[ bonds ]
; ai  aj funct   c0 c1  c2 c3
    1     2          1
    1    29          1
    1    30          1
    1    31          1
    2    17          1
    2     3          1
    2    21          1
    3    33          1
    3     4          1
    3    32          1
    4    34          1
    4     5          1
    4    35          1
    5    36          1
    5     6          1
    5    16          1
    6    13          1
    6     7          1
```

| | | |
|---|---|---|
| 6 | 8 | 1 |
| 7 | 37 | 1 |
| 7 | 39 | 1 |
| 7 | 38 | 1 |
| 8 | 9 | 1 |
| 8 | 41 | 1 |
| 8 | 40 | 1 |
| 9 | 10 | 1 |
| 9 | 43 | 1 |
| 9 | 42 | 1 |
| 10 | 44 | 1 |
| 10 | 11 | 1 |
| 10 | 12 | 1 |
| 11 | 45 | 1 |
| 12 | 13 | 1 |
| 12 | 46 | 1 |
| 12 | 47 | 1 |
| 13 | 48 | 1 |
| 13 | 14 | 1 |
| 14 | 49 | 1 |
| 14 | 50 | 1 |
| 14 | 15 | 1 |
| 15 | 51 | 1 |
| 15 | 52 | 1 |
| 15 | 16 | 1 |
| 16 | 17 | 1 |
| 16 | 53 | 1 |
| 17 | 19 | 1 |
| 17 | 18 | 1 |
| 18 | 54 | 1 |
| 19 | 20 | 1 |
| 19 | 55 | 1 |
| 19 | 56 | 1 |
| 20 | 57 | 1 |
| 20 | 58 | 1 |
| 20 | 21 | 1 |
| 21 | 59 | 1 |
| 21 | 22 | 1 |
| 22 | 28 | 1 |
| 22 | 23 | 1 |
| 23 | 60 | 1 |
| 23 | 24 | 1 |
| 24 | 25 | 1 |
| 24 | 61 | 1 |
| 25 | 26 | 1 |
| 25 | 27 | 1 |

```
   27    28             1
   28    62             1

[ pairs ]
; ai  aj funct   c0 c1  c2 c3
    1    33             1
    1     4             1
    1    16             1
    1    18             1
    1    19             1
    1    20             1
    1    22             1
    1    59             1
    1    32             1
    2    34             1
    2    35             1
    2     5             1
    2    23             1
    2    15             1
    2    53             1
    2    54             1
    2    55             1
    2    56             1
    2    57             1
    2    58             1
    2    28             1
    3    36             1
    3     6             1
    3    16             1
    3    18             1
    3    19             1
    3    20             1
    3    22             1
    3    59             1
    3    29             1
    3    30             1
    3    31             1
    4     7             1
    4     8             1
    4    13             1
    4    15             1
    4    17             1
    4    21             1
    4    53             1
    5    33             1
    5    19             1
```

| | | |
|---|---|---|
| 5 | 37 | 1 |
| 5 | 38 | 1 |
| 5 | 39 | 1 |
| 5 | 40 | 1 |
| 5 | 9 | 1 |
| 5 | 12 | 1 |
| 5 | 14 | 1 |
| 5 | 48 | 1 |
| 5 | 41 | 1 |
| 5 | 51 | 1 |
| 5 | 52 | 1 |
| 5 | 18 | 1 |
| 5 | 32 | 1 |
| 6 | 34 | 1 |
| 6 | 49 | 1 |
| 6 | 10 | 1 |
| 6 | 43 | 1 |
| 6 | 35 | 1 |
| 6 | 15 | 1 |
| 6 | 46 | 1 |
| 6 | 17 | 1 |
| 6 | 50 | 1 |
| 6 | 53 | 1 |
| 6 | 47 | 1 |
| 6 | 42 | 1 |
| 7 | 40 | 1 |
| 7 | 9 | 1 |
| 7 | 12 | 1 |
| 7 | 14 | 1 |
| 7 | 48 | 1 |
| 7 | 41 | 1 |
| 7 | 36 | 1 |
| 7 | 16 | 1 |
| 8 | 37 | 1 |
| 8 | 38 | 1 |
| 8 | 39 | 1 |
| 8 | 11 | 1 |
| 8 | 12 | 1 |
| 8 | 14 | 1 |
| 8 | 48 | 1 |
| 8 | 44 | 1 |
| 8 | 36 | 1 |
| 8 | 16 | 1 |
| 9 | 45 | 1 |
| 9 | 13 | 1 |
| 9 | 46 | 1 |

| | | |
|---|---|---|
| 9 | 47 | 1 |
| 10 | 41 | 1 |
| 10 | 48 | 1 |
| 10 | 14 | 1 |
| 10 | 40 | 1 |
| 11 | 42 | 1 |
| 11 | 43 | 1 |
| 11 | 13 | 1 |
| 11 | 46 | 1 |
| 11 | 47 | 1 |
| 12 | 42 | 1 |
| 12 | 43 | 1 |
| 12 | 45 | 1 |
| 12 | 15 | 1 |
| 12 | 49 | 1 |
| 12 | 50 | 1 |
| 13 | 52 | 1 |
| 13 | 37 | 1 |
| 13 | 38 | 1 |
| 13 | 39 | 1 |
| 13 | 40 | 1 |
| 13 | 44 | 1 |
| 13 | 16 | 1 |
| 13 | 41 | 1 |
| 13 | 51 | 1 |
| 13 | 36 | 1 |
| 14 | 46 | 1 |
| 14 | 47 | 1 |
| 14 | 17 | 1 |
| 14 | 53 | 1 |
| 15 | 48 | 1 |
| 15 | 18 | 1 |
| 15 | 19 | 1 |
| 15 | 36 | 1 |
| 16 | 34 | 1 |
| 16 | 35 | 1 |
| 16 | 49 | 1 |
| 16 | 50 | 1 |
| 16 | 20 | 1 |
| 16 | 21 | 1 |
| 16 | 54 | 1 |
| 16 | 55 | 1 |
| 16 | 56 | 1 |
| 17 | 33 | 1 |
| 17 | 52 | 1 |
| 17 | 51 | 1 |

| | | |
|---|---|---|
| 17 | 36 | 1 |
| 17 | 22 | 1 |
| 17 | 57 | 1 |
| 17 | 58 | 1 |
| 17 | 59 | 1 |
| 17 | 29 | 1 |
| 17 | 30 | 1 |
| 17 | 31 | 1 |
| 17 | 32 | 1 |
| 18 | 20 | 1 |
| 18 | 53 | 1 |
| 18 | 55 | 1 |
| 18 | 56 | 1 |
| 18 | 21 | 1 |
| 19 | 53 | 1 |
| 19 | 54 | 1 |
| 19 | 59 | 1 |
| 19 | 22 | 1 |
| 20 | 23 | 1 |
| 20 | 28 | 1 |
| 21 | 33 | 1 |
| 21 | 24 | 1 |
| 21 | 62 | 1 |
| 21 | 55 | 1 |
| 21 | 56 | 1 |
| 21 | 27 | 1 |
| 21 | 60 | 1 |
| 21 | 29 | 1 |
| 21 | 30 | 1 |
| 21 | 31 | 1 |
| 21 | 32 | 1 |
| 22 | 25 | 1 |
| 22 | 57 | 1 |
| 22 | 58 | 1 |
| 22 | 61 | 1 |
| 23 | 62 | 1 |
| 23 | 26 | 1 |
| 23 | 59 | 1 |
| 23 | 27 | 1 |
| 24 | 28 | 1 |
| 25 | 60 | 1 |
| 25 | 62 | 1 |
| 26 | 28 | 1 |
| 26 | 61 | 1 |
| 27 | 61 | 1 |
| 28 | 59 | 1 |

```
  28   60          1
  32   34          1
  32   35          1
  33   34          1
  33   35          1
  34   36          1
  35   36          1
  36   53          1
  40   42          1
  40   43          1
  41   42          1
  41   43          1
  42   44          1
  43   44          1
  44   45          1
  44   46          1
  44   47          1
  46   48          1
  47   48          1
  48   49          1
  48   50          1
  49   51          1
  49   52          1
  50   51          1
  50   52          1
  51   53          1
  52   53          1
  55   57          1
  55   58          1
  56   57          1
  56   58          1
  57   59          1
  58   59          1
  60   61          1

[ angles ]
; ai  aj ak funct  c0  c1 c2 c3
   2    1    29    5
   2    1    30    5
   2    1    31    5
  29     1    30   5
  29     1    31   5
  30     1    31   5
   1    2    17    5
   1    2     3    5
   1    2    21    5
```

| | | | |
|---|---|---|---|
| 17 | 2 | 3 | 5 |
| 17 | 2 | 21 | 5 |
| 3 | 2 | 21 | 5 |
| 33 | 3 | 2 | 5 |
| 33 | 3 | 4 | 5 |
| 33 | 3 | 32 | 5 |
| 2 | 3 | 4 | 5 |
| 2 | 3 | 32 | 5 |
| 4 | 3 | 32 | 5 |
| 34 | 4 | 3 | 5 |
| 34 | 4 | 5 | 5 |
| 34 | 4 | 35 | 5 |
| 3 | 4 | 5 | 5 |
| 3 | 4 | 35 | 5 |
| 5 | 4 | 35 | 5 |
| 36 | 5 | 4 | 5 |
| 36 | 5 | 6 | 5 |
| 36 | 5 | 16 | 5 |
| 4 | 5 | 6 | 5 |
| 4 | 5 | 16 | 5 |
| 6 | 5 | 16 | 5 |
| 13 | 6 | 5 | 5 |
| 13 | 6 | 7 | 5 |
| 13 | 6 | 8 | 5 |
| 5 | 6 | 7 | 5 |
| 5 | 6 | 8 | 5 |
| 7 | 6 | 8 | 5 |
| 37 | 7 | 6 | 5 |
| 37 | 7 | 39 | 5 |
| 37 | 7 | 38 | 5 |
| 6 | 7 | 39 | 5 |
| 6 | 7 | 38 | 5 |
| 39 | 7 | 38 | 5 |
| 9 | 8 | 41 | 5 |
| 9 | 8 | 6 | 5 |
| 9 | 8 | 40 | 5 |
| 41 | 8 | 6 | 5 |
| 41 | 8 | 40 | 5 |
| 6 | 8 | 40 | 5 |
| 10 | 9 | 43 | 5 |
| 10 | 9 | 42 | 5 |
| 10 | 9 | 8 | 5 |
| 43 | 9 | 42 | 5 |
| 43 | 9 | 8 | 5 |
| 42 | 9 | 8 | 5 |
| 9 | 10 | 44 | 5 |

| | | | |
|---|---|---|---|
| 9 | 10 | 11 | 5 |
| 9 | 10 | 12 | 5 |
| 44 | 10 | 11 | 5 |
| 44 | 10 | 12 | 5 |
| 11 | 10 | 12 | 5 |
| 10 | 11 | 45 | 5 |
| 10 | 12 | 13 | 5 |
| 10 | 12 | 46 | 5 |
| 10 | 12 | 47 | 5 |
| 13 | 12 | 46 | 5 |
| 13 | 12 | 47 | 5 |
| 46 | 12 | 47 | 5 |
| 48 | 13 | 12 | 5 |
| 48 | 13 | 6 | 5 |
| 48 | 13 | 14 | 5 |
| 12 | 13 | 6 | 5 |
| 12 | 13 | 14 | 5 |
| 6 | 13 | 14 | 5 |
| 49 | 14 | 50 | 5 |
| 49 | 14 | 13 | 5 |
| 49 | 14 | 15 | 5 |
| 50 | 14 | 13 | 5 |
| 50 | 14 | 15 | 5 |
| 13 | 14 | 15 | 5 |
| 51 | 15 | 52 | 5 |
| 51 | 15 | 14 | 5 |
| 51 | 15 | 16 | 5 |
| 52 | 15 | 14 | 5 |
| 52 | 15 | 16 | 5 |
| 14 | 15 | 16 | 5 |
| 17 | 16 | 53 | 5 |
| 17 | 16 | 5 | 5 |
| 17 | 16 | 15 | 5 |
| 53 | 16 | 5 | 5 |
| 53 | 16 | 15 | 5 |
| 5 | 16 | 15 | 5 |
| 2 | 17 | 19 | 5 |
| 2 | 17 | 18 | 5 |
| 2 | 17 | 16 | 5 |
| 19 | 17 | 18 | 5 |
| 19 | 17 | 16 | 5 |
| 18 | 17 | 16 | 5 |
| 17 | 18 | 54 | 5 |
| 17 | 19 | 20 | 5 |
| 17 | 19 | 55 | 5 |
| 17 | 19 | 56 | 5 |

```
   20    19    55   5
   20    19    56   5
   55    19    56   5
   57    20    58   5
   57    20    19   5
   57    20    21   5
   58    20    19   5
   58    20    21   5
   19    20    21   5
    2    21    59   5
    2    21    20   5
    2    21    22   5
   59    21    20   5
   59    21    22   5
   20    21    22   5
   28    22    21   5
   28    22    23   5
   21    22    23   5
   60    23    22   5
   60    23    24   5
   22    23    24   5
   25    24    61   5
   25    24    23   5
   61    24    23   5
   26    25    27   5
   26    25    24   5
   27    25    24   5
   25    27    28   5
   27    28    22   5
   27    28    62   5
   22    28    62   5

[ dihedrals ]
; ai  aj ak  al funct   c0 c1 c2 c3 c4 c5
   29     1     2    17                    9
   29     1     2     3                    9
   29     1     2    21                    9
   30     1     2    17                    9
   30     1     2     3                    9
   30     1     2    21                    9
   31     1     2    17                    9
   31     1     2     3                    9
   31     1     2    21                    9
    1     2    17    19                    9
    1     2    17    18                    9
    1     2    17    16                    9
```

| | | | | |
|---|---|---|---|---|
| 3 | 2 | 17 | 19 | 9 |
| 3 | 2 | 17 | 18 | 9 |
| 3 | 2 | 17 | 16 | 9 |
| 21 | 2 | 17 | 19 | 9 |
| 21 | 2 | 17 | 18 | 9 |
| 21 | 2 | 17 | 16 | 9 |
| 1 | 2 | 3 | 33 | 9 |
| 1 | 2 | 3 | 4 | 9 |
| 1 | 2 | 3 | 32 | 9 |
| 17 | 2 | 3 | 33 | 9 |
| 17 | 2 | 3 | 4 | 9 |
| 17 | 2 | 3 | 32 | 9 |
| 21 | 2 | 3 | 33 | 9 |
| 21 | 2 | 3 | 4 | 9 |
| 21 | 2 | 3 | 32 | 9 |
| 1 | 2 | 21 | 59 | 9 |
| 1 | 2 | 21 | 20 | 9 |
| 1 | 2 | 21 | 22 | 9 |
| 17 | 2 | 21 | 59 | 9 |
| 17 | 2 | 21 | 20 | 9 |
| 17 | 2 | 21 | 22 | 9 |
| 3 | 2 | 21 | 59 | 9 |
| 3 | 2 | 21 | 20 | 9 |
| 3 | 2 | 21 | 22 | 9 |
| 33 | 3 | 4 | 34 | 9 |
| 33 | 3 | 4 | 5 | 9 |
| 33 | 3 | 4 | 35 | 9 |
| 2 | 3 | 4 | 34 | 9 |
| 2 | 3 | 4 | 5 | 9 |
| 2 | 3 | 4 | 35 | 9 |
| 32 | 3 | 4 | 34 | 9 |
| 32 | 3 | 4 | 5 | 9 |
| 32 | 3 | 4 | 35 | 9 |
| 34 | 4 | 5 | 36 | 9 |
| 34 | 4 | 5 | 6 | 9 |
| 34 | 4 | 5 | 16 | 9 |
| 3 | 4 | 5 | 36 | 9 |
| 3 | 4 | 5 | 6 | 9 |
| 3 | 4 | 5 | 16 | 9 |
| 35 | 4 | 5 | 36 | 9 |
| 35 | 4 | 5 | 6 | 9 |
| 35 | 4 | 5 | 16 | 9 |
| 36 | 5 | 6 | 13 | 9 |
| 36 | 5 | 6 | 7 | 9 |
| 36 | 5 | 6 | 8 | 9 |
| 4 | 5 | 6 | 13 | 9 |

| | | | | |
|---|---|---|---|---|
| 4 | 5 | 6 | 7 | 9 |
| 4 | 5 | 6 | 8 | 9 |
| 16 | 5 | 6 | 13 | 9 |
| 16 | 5 | 6 | 7 | 9 |
| 16 | 5 | 6 | 8 | 9 |
| 36 | 5 | 16 | 17 | 9 |
| 36 | 5 | 16 | 53 | 9 |
| 36 | 5 | 16 | 15 | 9 |
| 4 | 5 | 16 | 17 | 9 |
| 4 | 5 | 16 | 53 | 9 |
| 4 | 5 | 16 | 15 | 9 |
| 6 | 5 | 16 | 17 | 9 |
| 6 | 5 | 16 | 53 | 9 |
| 6 | 5 | 16 | 15 | 9 |
| 5 | 6 | 13 | 48 | 9 |
| 5 | 6 | 13 | 12 | 9 |
| 5 | 6 | 13 | 14 | 9 |
| 7 | 6 | 13 | 48 | 9 |
| 7 | 6 | 13 | 12 | 9 |
| 7 | 6 | 13 | 14 | 9 |
| 8 | 6 | 13 | 48 | 9 |
| 8 | 6 | 13 | 12 | 9 |
| 8 | 6 | 13 | 14 | 9 |
| 13 | 6 | 7 | 37 | 9 |
| 13 | 6 | 7 | 39 | 9 |
| 13 | 6 | 7 | 38 | 9 |
| 5 | 6 | 7 | 37 | 9 |
| 5 | 6 | 7 | 39 | 9 |
| 5 | 6 | 7 | 38 | 9 |
| 8 | 6 | 7 | 37 | 9 |
| 8 | 6 | 7 | 39 | 9 |
| 8 | 6 | 7 | 38 | 9 |
| 13 | 6 | 8 | 9 | 9 |
| 13 | 6 | 8 | 41 | 9 |
| 13 | 6 | 8 | 40 | 9 |
| 5 | 6 | 8 | 9 | 9 |
| 5 | 6 | 8 | 41 | 9 |
| 5 | 6 | 8 | 40 | 9 |
| 7 | 6 | 8 | 9 | 9 |
| 7 | 6 | 8 | 41 | 9 |
| 7 | 6 | 8 | 40 | 9 |
| 41 | 8 | 9 | 10 | 9 |
| 41 | 8 | 9 | 43 | 9 |
| 41 | 8 | 9 | 42 | 9 |
| 6 | 8 | 9 | 10 | 9 |
| 6 | 8 | 9 | 43 | 9 |

| | | | | |
|---|---|---|---|---|
| 6 | 8 | 9 | 42 | 9 |
| 40 | 8 | 9 | 10 | 9 |
| 40 | 8 | 9 | 43 | 9 |
| 40 | 8 | 9 | 42 | 9 |
| 43 | 9 | 10 | 44 | 9 |
| 43 | 9 | 10 | 11 | 9 |
| 43 | 9 | 10 | 12 | 9 |
| 42 | 9 | 10 | 44 | 9 |
| 42 | 9 | 10 | 11 | 9 |
| 42 | 9 | 10 | 12 | 9 |
| 8 | 9 | 10 | 44 | 9 |
| 8 | 9 | 10 | 11 | 9 |
| 8 | 9 | 10 | 12 | 9 |
| 9 | 10 | 11 | 45 | 9 |
| 44 | 10 | 11 | 45 | 9 |
| 12 | 10 | 11 | 45 | 9 |
| 9 | 10 | 12 | 13 | 9 |
| 9 | 10 | 12 | 46 | 9 |
| 9 | 10 | 12 | 47 | 9 |
| 44 | 10 | 12 | 13 | 9 |
| 44 | 10 | 12 | 46 | 9 |
| 44 | 10 | 12 | 47 | 9 |
| 11 | 10 | 12 | 13 | 9 |
| 11 | 10 | 12 | 46 | 9 |
| 11 | 10 | 12 | 47 | 9 |
| 10 | 12 | 13 | 48 | 9 |
| 10 | 12 | 13 | 6 | 9 |
| 10 | 12 | 13 | 14 | 9 |
| 46 | 12 | 13 | 48 | 9 |
| 46 | 12 | 13 | 6 | 9 |
| 46 | 12 | 13 | 14 | 9 |
| 47 | 12 | 13 | 48 | 9 |
| 47 | 12 | 13 | 6 | 9 |
| 47 | 12 | 13 | 14 | 9 |
| 48 | 13 | 14 | 49 | 9 |
| 48 | 13 | 14 | 50 | 9 |
| 48 | 13 | 14 | 15 | 9 |
| 12 | 13 | 14 | 49 | 9 |
| 12 | 13 | 14 | 50 | 9 |
| 12 | 13 | 14 | 15 | 9 |
| 6 | 13 | 14 | 49 | 9 |
| 6 | 13 | 14 | 50 | 9 |
| 6 | 13 | 14 | 15 | 9 |
| 49 | 14 | 15 | 51 | 9 |
| 49 | 14 | 15 | 52 | 9 |
| 49 | 14 | 15 | 16 | 9 |

| | | | | |
|---|---|---|---|---|
| 50 | 14 | 15 | 51 | 9 |
| 50 | 14 | 15 | 52 | 9 |
| 50 | 14 | 15 | 16 | 9 |
| 13 | 14 | 15 | 51 | 9 |
| 13 | 14 | 15 | 52 | 9 |
| 13 | 14 | 15 | 16 | 9 |
| 51 | 15 | 16 | 17 | 9 |
| 51 | 15 | 16 | 53 | 9 |
| 51 | 15 | 16 | 5 | 9 |
| 52 | 15 | 16 | 17 | 9 |
| 52 | 15 | 16 | 53 | 9 |
| 52 | 15 | 16 | 5 | 9 |
| 14 | 15 | 16 | 17 | 9 |
| 14 | 15 | 16 | 53 | 9 |
| 14 | 15 | 16 | 5 | 9 |
| 53 | 16 | 17 | 2 | 9 |
| 53 | 16 | 17 | 19 | 9 |
| 53 | 16 | 17 | 18 | 9 |
| 5 | 16 | 17 | 2 | 9 |
| 5 | 16 | 17 | 19 | 9 |
| 5 | 16 | 17 | 18 | 9 |
| 15 | 16 | 17 | 2 | 9 |
| 15 | 16 | 17 | 19 | 9 |
| 15 | 16 | 17 | 18 | 9 |
| 2 | 17 | 19 | 20 | 9 |
| 2 | 17 | 19 | 55 | 9 |
| 2 | 17 | 19 | 56 | 9 |
| 18 | 17 | 19 | 20 | 9 |
| 18 | 17 | 19 | 55 | 9 |
| 18 | 17 | 19 | 56 | 9 |
| 16 | 17 | 19 | 20 | 9 |
| 16 | 17 | 19 | 55 | 9 |
| 16 | 17 | 19 | 56 | 9 |
| 2 | 17 | 18 | 54 | 9 |
| 19 | 17 | 18 | 54 | 9 |
| 16 | 17 | 18 | 54 | 9 |
| 17 | 19 | 20 | 57 | 9 |
| 17 | 19 | 20 | 58 | 9 |
| 17 | 19 | 20 | 21 | 9 |
| 55 | 19 | 20 | 57 | 9 |
| 55 | 19 | 20 | 58 | 9 |
| 55 | 19 | 20 | 21 | 9 |
| 56 | 19 | 20 | 57 | 9 |
| 56 | 19 | 20 | 58 | 9 |
| 56 | 19 | 20 | 21 | 9 |
| 57 | 20 | 21 | 2 | 9 |

```
   57    20    21    59                      9
   57    20    21    22                      9
   58    20    21     2                      9
   58    20    21    59                      9
   58    20    21    22                      9
   19    20    21     2                      9
   19    20    21    59                      9
   19    20    21    22                      9
    2    21    22    28                      9
    2    21    22    23                      9
   59    21    22    28                      9
   59    21    22    23                      9
   20    21    22    28                      9
   20    21    22    23                      9
   21    22    28    27                      9
   21    22    28    62                      9
   23    22    28    27                      9
   23    22    28    62                      9
   28    22    23    60                      9
   28    22    23    24                      9
   21    22    23    60                      9
   21    22    23    24                      9
   60    23    24    25                      9
   60    23    24    61                      9
   22    23    24    25                      9
   22    23    24    61                      9
   61    24    25    26                      9
   61    24    25    27                      9
   23    24    25    26                      9
   23    24    25    27                      9
   26    25    27    28                      9
   24    25    27    28                      9
   25    27    28    22                      9
   25    27    28    62                      9

[ dihedrals ]
; ai  aj ak al funct c0 c1  c2 c3
   25    24    26    27                      2

; Include Position restraint file
#ifdef POSRES
#include "bufalin_posres.itp"
#endif
```

**Supporting data file S2.** Bufalin prm file.

```
[ bondtypes ]
;      i        j  func           b0          kb
  CG2R62   CG3C51     1   0.14900000    192464.00
  CG3RC1    OG311     1   0.14200000    358150.40

[ angletypes ]
;      i        j        k  func       theta0       ktheta          ub0          kub
  CG2R62   CG2R62   CG3C51     5   124.200000   334.720000   0.00000000         0.00
  OG3R60   CG2R62    HGR62     5   119.000000   292.880000   0.00000000         0.00
  CG2R62   CG3C51   CG3C52     5   112.300000   435.136000   0.00000000         0.00
  CG2R62   CG3C51   CG3RC1     5   112.300000   435.136000   0.00000000         0.00
  CG2R62   CG3C51     HGA1     5   112.000000   418.400000   0.00000000         0.00
   CG311   CG3RC1    OG311     5   113.500000   488.272800   0.25610000      9338.69
  CG3C52   CG3RC1    OG311     5   110.000000   633.457600   0.00000000         0.00
  CG3RC1   CG3RC1    OG311     5   111.000000   446.432800   0.25610000      6694.40
  CG3RC1    OG311     HGP1     5   109.000000   418.400000   0.00000000         0.00

[ dihedraltypes ]
;      i        j        k        l  func         phi0         kphi  mult
  CG2R62   CG2R62   CG2R62   CG3C51     9   180.000000    12.970400     2
  CG3C51   CG2R62   CG2R62   OG3R60     9   180.000000    10.041600     2
  CG3C51   CG2R62   CG2R62    HGR62     9   180.000000    16.736000     2
  CG2R62   CG2R62   CG3C51   CG3C52     9   180.000000     0.962320     2
  CG2R62   CG2R62   CG3C51   CG3RC1     9   180.000000     0.962320     2
  CG2R62   CG2R62   CG3C51     HGA1     9     0.000000     0.008368     6
   HGR62   CG2R62   OG3R60   CG2R63     9     0.000000     3.179840     2
   CG311    CG311   CG3RC1    OG311     9     0.000000     0.209200     3
   CG321    CG311   CG3RC1    OG311     9     0.000000     0.209200     3
    HGA1    CG311   CG3RC1    OG311     9     0.000000     0.209200     3
  CG2R62   CG3C51   CG3C52   CG3C52     9     0.000000     0.585760     3
  CG2R62   CG3C51   CG3C52     HGA2     9     0.000000     0.585760     3
  CG2R62   CG3C51   CG3RC1    CG321     9   180.000000     2.092000     2
  CG2R62   CG3C51   CG3RC1    CG331     9   180.000000     2.092000     2
  CG2R62   CG3C51   CG3RC1   CG3RC1     9     0.000000     0.627600     3
  CG3C52   CG3C52   CG3RC1    OG311     9   180.000000     2.092000     1
  CG3C52   CG3C52   CG3RC1    OG311     9     0.000000     2.928800     2
  CG3C52   CG3C52   CG3RC1    OG311     9     0.000000     1.673600     3
  CG3C52   CG3C52   CG3RC1    OG311     9     0.000000     1.673600     5
    HGA2   CG3C52   CG3RC1    OG311     9   180.000000     0.815880     3
   CG321   CG3RC1   CG3RC1    OG311     9     0.000000     0.627600     3
   CG331   CG3RC1   CG3RC1    OG311     9     0.000000     0.209200     3
  CG3C51   CG3RC1   CG3RC1    OG311     9     0.000000     0.209200     3
   CG311   CG3RC1    OG311     HGP1     9     0.000000     1.213360     1
   CG311   CG3RC1    OG311     HGP1     9     0.000000     2.594080     2
   CG311   CG3RC1    OG311     HGP1     9     0.000000     0.209200     3
```

```
  CG3C52   CG3RC1    OG311     HGP1     9     0.000000     1.213360     1
  CG3C52   CG3RC1    OG311     HGP1     9     0.000000     2.594080     2
  CG3C52   CG3RC1    OG311     HGP1     9     0.000000     0.209200     3
  CG3RC1   CG3RC1    OG311     HGP1     9     0.000000     6.276000     1
  CG3RC1   CG3RC1    OG311     HGP1     9   180.000000     1.255200     2
  CG3RC1   CG3RC1    OG311     HGP1     9     0.000000     1.338880     3

[ dihedraltypes ]
; 'improper' dihedrals
;      i        j        k        l  func         phi0         kphi
```

**Supporting data file S3.** Cinobufagin itp file.

```
[ moleculetype ]
; Name   nrexcl
cinobufagin        3

[ atoms ]
; nr   type resnr residue  atom   cgnr  charge mass  typeB    chargeB massB
; residue 1 cinobufagin rtp cinobufagin q            qsum
     1      CG331      1 cinobufagin    C18      1     -0.269     12.011   ;
     2     CG3RC1      1 cinobufagin    C13      2      0.003     12.011   ;
     3      CG321      1 cinobufagin    C12      3     -0.183     12.011   ;
     4      CG321      1 cinobufagin    C11      4     -0.182     12.011   ;
     5      CG311      1 cinobufagin     C9      5     -0.091     12.011   ;
     6      CG301      1 cinobufagin    C10      6     -0.001     12.011   ;
     7      CG331      1 cinobufagin    C19      7     -0.274     12.011   ;
     8      CG321      1 cinobufagin     C1      8     -0.180     12.011   ;
     9      CG321      1 cinobufagin     C2      9     -0.179     12.011   ;
    10      CG311      1 cinobufagin     C3     10      0.135     12.011   ;
    11      OG311      1 cinobufagin     O3     11     -0.650     15.999   ;
    12      CG321      1 cinobufagin     C4     12     -0.177     12.011   ;
    13      CG311      1 cinobufagin     C5     13     -0.090     12.011   ;
    14      CG321      1 cinobufagin     C6     14     -0.181     12.011   ;
    15      CG321      1 cinobufagin     C7     15     -0.182     12.011   ;
    16      CG311      1 cinobufagin     C8     16     -0.086     12.011   ;
    17     CG3C51      1 cinobufagin    C16     17      0.051     12.011   ;
    18     CG3C51      1 cinobufagin    C17     18     -0.033     12.011   ;
    19     CG2R62      1 cinobufagin    C20     19      0.031     12.011   ;
    20     CG2R62      1 cinobufagin    C22     20     -0.272     12.011   ;
    21     CG2R62      1 cinobufagin    C23     21     -0.347     12.011   ;
    22     CG2R63      1 cinobufagin    C24     22      0.490     12.011   ;
    23      OG2D4      1 cinobufagin    O24     23     -0.460     15.999   ;
    24     OG3R60      1 cinobufagin    O21     24     -0.362     15.999   ;
    25     CG2R62      1 cinobufagin    C21     25      0.119     12.011   ;
    26       HGA3      1 cinobufagin   H181     26      0.090      1.008   ;
    27       HGA3      1 cinobufagin   H182     27      0.090      1.008   ;
    28       HGA3      1 cinobufagin   H183     28      0.090      1.008   ;
    29       HGA2      1 cinobufagin   H121     29      0.090      1.008   ;
    30       HGA2      1 cinobufagin   H122     30      0.090      1.008   ;
    31       HGA2      1 cinobufagin   H111     31      0.090      1.008   ;
    32       HGA2      1 cinobufagin   H112     32      0.090      1.008   ;
    33       HGA1      1 cinobufagin     H9     33      0.090      1.008   ;
    34       HGA3      1 cinobufagin   H191     34      0.090      1.008   ;
    35       HGA3      1 cinobufagin   H192     35      0.090      1.008   ;
    36       HGA3      1 cinobufagin   H193     36      0.090      1.008   ;
    37       HGA2      1 cinobufagin    H11     37      0.090      1.008   ;
    38       HGA2      1 cinobufagin    H12     38      0.090      1.008   ;
```

```
    39       HGA2      1 cinobufagin    H21     39      0.090       1.008   ;
    40       HGA2      1 cinobufagin    H22     40      0.090       1.008   ;
    41       HGA1      1 cinobufagin     H3     41      0.090       1.008   ;
    42       HGP1      1 cinobufagin    HO3     42      0.419       1.008   ;
    43       HGA2      1 cinobufagin    H41     43      0.090       1.008   ;
    44       HGA2      1 cinobufagin    H42     44      0.090       1.008   ;
    45       HGA1      1 cinobufagin     H5     45      0.090       1.008   ;
    46       HGA2      1 cinobufagin    H61     46      0.090       1.008   ;
    47       HGA2      1 cinobufagin    H62     47      0.090       1.008   ;
    48       HGA2      1 cinobufagin    H71     48      0.090       1.008   ;
    49       HGA2      1 cinobufagin    H72     49      0.090       1.008   ;
    50       HGA1      1 cinobufagin     H8     50      0.090       1.008   ;
    51       HGA1      1 cinobufagin   H161     51      0.090       1.008   ;
    52      OG302      1 cinobufagin    O52     52     -0.370      15.999   ;
    53       HGA1      1 cinobufagin    H17     53      0.090       1.008   ;
    54      HGR62      1 cinobufagin     H2     54      0.268       1.008   ;
    55      HGR62      1 cinobufagin    H23     55      0.199       1.008   ;
    56      HGR62      1 cinobufagin     H1     56      0.284       1.008   ;
    57     CG3RC1      1 cinobufagin    C57     57      0.191      12.011   ;
    58     CG3RC1      1 cinobufagin    C58     58      0.124      12.011   ;
    59     OG3C31      1 cinobufagin    O59     59     -0.404      15.999   ;
    60       HGA1      1 cinobufagin    H60     60      0.090       1.008   ;
    61      CG2O2      1 cinobufagin    C61     61      0.897      12.011   ;
    62      CG331      1 cinobufagin    C62     62     -0.311      12.011   ;
    63      OG2D1      1 cinobufagin    O63     63     -0.627      15.999   ;
    64       HGA3      1 cinobufagin    H64     64      0.090       1.008   ;
    65       HGA3      1 cinobufagin    H65     65      0.090       1.008   ;
    66       HGA3      1 cinobufagin    H66     66      0.090       1.008   ;

[ bonds ]
; ai  aj funct   c0 c1  c2 c3
    1     2       1
    1    27       1
    1    28       1
    1    26       1
    2    18       1
    2     3       1
    2    57       1
    3     4       1
    3    29       1
    3    30       1
    4     5       1
    4    31       1
    4    32       1
    5    33       1
    5     6       1
```

| | | |
|---|---|---|
| 5 | 16 | 1 |
| 6 | 13 | 1 |
| 6 | 7 | 1 |
| 6 | 8 | 1 |
| 7 | 34 | 1 |
| 7 | 35 | 1 |
| 7 | 36 | 1 |
| 8 | 9 | 1 |
| 8 | 37 | 1 |
| 8 | 38 | 1 |
| 9 | 10 | 1 |
| 9 | 40 | 1 |
| 9 | 39 | 1 |
| 10 | 41 | 1 |
| 10 | 11 | 1 |
| 10 | 12 | 1 |
| 11 | 42 | 1 |
| 12 | 43 | 1 |
| 12 | 44 | 1 |
| 12 | 13 | 1 |
| 13 | 45 | 1 |
| 13 | 14 | 1 |
| 14 | 47 | 1 |
| 14 | 46 | 1 |
| 14 | 15 | 1 |
| 15 | 49 | 1 |
| 15 | 48 | 1 |
| 15 | 16 | 1 |
| 16 | 57 | 1 |
| 16 | 50 | 1 |
| 17 | 18 | 1 |
| 17 | 51 | 1 |
| 17 | 52 | 1 |
| 17 | 58 | 1 |
| 18 | 19 | 1 |
| 18 | 53 | 1 |
| 19 | 25 | 1 |
| 19 | 20 | 1 |
| 20 | 21 | 1 |
| 20 | 54 | 1 |
| 21 | 22 | 1 |
| 21 | 55 | 1 |
| 22 | 23 | 1 |
| 22 | 24 | 1 |
| 24 | 25 | 1 |
| 25 | 56 | 1 |

```
  52   61          1
  57   59          1
  57   58          1
  58   59          1
  58   60          1
  61   62          1
  61   63          1
  62   65          1
  62   66          1
  62   64          1

[ pairs ]
; ai  aj funct   c0 c1 c2 c3
   1    4          1
   1   16          1
   1   17          1
   1   19          1
   1   53          1
   1   58          1
   1   59          1
   1   29          1
   1   30          1
   2   20          1
   2    5          1
   2   15          1
   2   50          1
   2   51          1
   2   52          1
   2   25          1
   2   60          1
   2   31          1
   2   32          1
   3   33          1
   3    6          1
   3   16          1
   3   17          1
   3   19          1
   3   53          1
   3   58          1
   3   26          1
   3   27          1
   3   28          1
   3   59          1
   4   50          1
   4    7          1
   4    8          1
```

| | | |
|---|---|---|
| 4 | 13 | 1 |
| 4 | 15 | 1 |
| 4 | 18 | 1 |
| 4 | 57 | 1 |
| 5 | 35 | 1 |
| 5 | 36 | 1 |
| 5 | 37 | 1 |
| 5 | 38 | 1 |
| 5 | 34 | 1 |
| 5 | 9 | 1 |
| 5 | 12 | 1 |
| 5 | 45 | 1 |
| 5 | 14 | 1 |
| 5 | 48 | 1 |
| 5 | 49 | 1 |
| 5 | 58 | 1 |
| 5 | 59 | 1 |
| 5 | 29 | 1 |
| 5 | 30 | 1 |
| 6 | 39 | 1 |
| 6 | 40 | 1 |
| 6 | 10 | 1 |
| 6 | 43 | 1 |
| 6 | 44 | 1 |
| 6 | 46 | 1 |
| 6 | 15 | 1 |
| 6 | 50 | 1 |
| 6 | 47 | 1 |
| 6 | 57 | 1 |
| 6 | 31 | 1 |
| 6 | 32 | 1 |
| 7 | 33 | 1 |
| 7 | 37 | 1 |
| 7 | 38 | 1 |
| 7 | 9 | 1 |
| 7 | 12 | 1 |
| 7 | 45 | 1 |
| 7 | 14 | 1 |
| 7 | 16 | 1 |
| 8 | 33 | 1 |
| 8 | 34 | 1 |
| 8 | 35 | 1 |
| 8 | 41 | 1 |
| 8 | 11 | 1 |
| 8 | 12 | 1 |
| 8 | 45 | 1 |

| | | |
|---|---|---|
| 8 | 14 | 1 |
| 8 | 16 | 1 |
| 8 | 36 | 1 |
| 9 | 42 | 1 |
| 9 | 43 | 1 |
| 9 | 44 | 1 |
| 9 | 13 | 1 |
| 10 | 45 | 1 |
| 10 | 14 | 1 |
| 10 | 37 | 1 |
| 10 | 38 | 1 |
| 11 | 39 | 1 |
| 11 | 43 | 1 |
| 11 | 44 | 1 |
| 11 | 13 | 1 |
| 11 | 40 | 1 |
| 12 | 39 | 1 |
| 12 | 40 | 1 |
| 12 | 42 | 1 |
| 12 | 46 | 1 |
| 12 | 47 | 1 |
| 12 | 15 | 1 |
| 13 | 33 | 1 |
| 13 | 34 | 1 |
| 13 | 35 | 1 |
| 13 | 37 | 1 |
| 13 | 38 | 1 |
| 13 | 16 | 1 |
| 13 | 49 | 1 |
| 13 | 41 | 1 |
| 13 | 36 | 1 |
| 13 | 48 | 1 |
| 14 | 43 | 1 |
| 14 | 44 | 1 |
| 14 | 50 | 1 |
| 14 | 57 | 1 |
| 15 | 33 | 1 |
| 15 | 45 | 1 |
| 15 | 58 | 1 |
| 15 | 59 | 1 |
| 16 | 46 | 1 |
| 16 | 47 | 1 |
| 16 | 17 | 1 |
| 16 | 18 | 1 |
| 16 | 60 | 1 |
| 16 | 31 | 1 |

| | | |
|---|---|---|
| 16 | 32 | 1 |
| 17 | 20 | 1 |
| 17 | 25 | 1 |
| 17 | 62 | 1 |
| 17 | 63 | 1 |
| 18 | 61 | 1 |
| 18 | 24 | 1 |
| 18 | 60 | 1 |
| 18 | 59 | 1 |
| 18 | 21 | 1 |
| 18 | 54 | 1 |
| 18 | 56 | 1 |
| 18 | 26 | 1 |
| 18 | 27 | 1 |
| 18 | 28 | 1 |
| 18 | 29 | 1 |
| 18 | 30 | 1 |
| 19 | 51 | 1 |
| 19 | 52 | 1 |
| 19 | 22 | 1 |
| 19 | 55 | 1 |
| 19 | 57 | 1 |
| 19 | 58 | 1 |
| 20 | 56 | 1 |
| 20 | 53 | 1 |
| 20 | 23 | 1 |
| 20 | 24 | 1 |
| 21 | 25 | 1 |
| 22 | 54 | 1 |
| 22 | 56 | 1 |
| 23 | 25 | 1 |
| 23 | 55 | 1 |
| 24 | 55 | 1 |
| 25 | 53 | 1 |
| 25 | 54 | 1 |
| 26 | 57 | 1 |
| 27 | 57 | 1 |
| 28 | 57 | 1 |
| 29 | 57 | 1 |
| 29 | 31 | 1 |
| 29 | 32 | 1 |
| 30 | 57 | 1 |
| 30 | 31 | 1 |
| 30 | 32 | 1 |
| 31 | 33 | 1 |
| 32 | 33 | 1 |

| | | |
|---|---|---|
| 33 | 50 | 1 |
| 33 | 57 | 1 |
| 37 | 39 | 1 |
| 37 | 40 | 1 |
| 38 | 39 | 1 |
| 38 | 40 | 1 |
| 39 | 41 | 1 |
| 40 | 41 | 1 |
| 41 | 42 | 1 |
| 41 | 43 | 1 |
| 41 | 44 | 1 |
| 43 | 45 | 1 |
| 44 | 45 | 1 |
| 45 | 46 | 1 |
| 45 | 47 | 1 |
| 46 | 49 | 1 |
| 46 | 48 | 1 |
| 47 | 49 | 1 |
| 47 | 48 | 1 |
| 48 | 50 | 1 |
| 48 | 57 | 1 |
| 49 | 50 | 1 |
| 49 | 57 | 1 |
| 50 | 58 | 1 |
| 50 | 59 | 1 |
| 51 | 53 | 1 |
| 51 | 57 | 1 |
| 51 | 59 | 1 |
| 51 | 60 | 1 |
| 51 | 61 | 1 |
| 52 | 65 | 1 |
| 52 | 66 | 1 |
| 52 | 53 | 1 |
| 52 | 57 | 1 |
| 52 | 59 | 1 |
| 52 | 60 | 1 |
| 52 | 64 | 1 |
| 53 | 57 | 1 |
| 53 | 58 | 1 |
| 54 | 55 | 1 |
| 58 | 61 | 1 |
| 63 | 65 | 1 |
| 63 | 64 | 1 |
| 63 | 66 | 1 |

[ angles ]

| ; ai | aj | ak | funct | c0 | c1 | c2 | c3 |
|---|---|---|---|---|---|---|---|
| 2 | 1 | 27 | 5 | | | | |
| 2 | 1 | 28 | 5 | | | | |
| 2 | 1 | 26 | 5 | | | | |
| 27 | 1 | 28 | 5 | | | | |
| 27 | 1 | 26 | 5 | | | | |
| 28 | 1 | 26 | 5 | | | | |
| 1 | 2 | 18 | 5 | | | | |
| 1 | 2 | 3 | 5 | | | | |
| 1 | 2 | 57 | 5 | | | | |
| 18 | 2 | 3 | 5 | | | | |
| 18 | 2 | 57 | 5 | | | | |
| 3 | 2 | 57 | 5 | | | | |
| 2 | 3 | 4 | 5 | | | | |
| 2 | 3 | 29 | 5 | | | | |
| 2 | 3 | 30 | 5 | | | | |
| 4 | 3 | 29 | 5 | | | | |
| 4 | 3 | 30 | 5 | | | | |
| 29 | 3 | 30 | 5 | | | | |
| 3 | 4 | 5 | 5 | | | | |
| 3 | 4 | 31 | 5 | | | | |
| 3 | 4 | 32 | 5 | | | | |
| 5 | 4 | 31 | 5 | | | | |
| 5 | 4 | 32 | 5 | | | | |
| 31 | 4 | 32 | 5 | | | | |
| 33 | 5 | 4 | 5 | | | | |
| 33 | 5 | 6 | 5 | | | | |
| 33 | 5 | 16 | 5 | | | | |
| 4 | 5 | 6 | 5 | | | | |
| 4 | 5 | 16 | 5 | | | | |
| 6 | 5 | 16 | 5 | | | | |
| 13 | 6 | 5 | 5 | | | | |
| 13 | 6 | 7 | 5 | | | | |
| 13 | 6 | 8 | 5 | | | | |
| 5 | 6 | 7 | 5 | | | | |
| 5 | 6 | 8 | 5 | | | | |
| 7 | 6 | 8 | 5 | | | | |
| 34 | 7 | 35 | 5 | | | | |
| 34 | 7 | 36 | 5 | | | | |
| 34 | 7 | 6 | 5 | | | | |
| 35 | 7 | 36 | 5 | | | | |
| 35 | 7 | 6 | 5 | | | | |
| 36 | 7 | 6 | 5 | | | | |
| 9 | 8 | 37 | 5 | | | | |
| 9 | 8 | 6 | 5 | | | | |
| 9 | 8 | 38 | 5 | | | | |

| | | | |
|---|---|---|---|
| 37 | 8 | 6 | 5 |
| 37 | 8 | 38 | 5 |
| 6 | 8 | 38 | 5 |
| 10 | 9 | 40 | 5 |
| 10 | 9 | 39 | 5 |
| 10 | 9 | 8 | 5 |
| 40 | 9 | 39 | 5 |
| 40 | 9 | 8 | 5 |
| 39 | 9 | 8 | 5 |
| 9 | 10 | 41 | 5 |
| 9 | 10 | 11 | 5 |
| 9 | 10 | 12 | 5 |
| 41 | 10 | 11 | 5 |
| 41 | 10 | 12 | 5 |
| 11 | 10 | 12 | 5 |
| 10 | 11 | 42 | 5 |
| 10 | 12 | 43 | 5 |
| 10 | 12 | 44 | 5 |
| 10 | 12 | 13 | 5 |
| 43 | 12 | 44 | 5 |
| 43 | 12 | 13 | 5 |
| 44 | 12 | 13 | 5 |
| 12 | 13 | 45 | 5 |
| 12 | 13 | 6 | 5 |
| 12 | 13 | 14 | 5 |
| 45 | 13 | 6 | 5 |
| 45 | 13 | 14 | 5 |
| 6 | 13 | 14 | 5 |
| 47 | 14 | 13 | 5 |
| 47 | 14 | 46 | 5 |
| 47 | 14 | 15 | 5 |
| 13 | 14 | 46 | 5 |
| 13 | 14 | 15 | 5 |
| 46 | 14 | 15 | 5 |
| 49 | 15 | 48 | 5 |
| 49 | 15 | 14 | 5 |
| 49 | 15 | 16 | 5 |
| 48 | 15 | 14 | 5 |
| 48 | 15 | 16 | 5 |
| 14 | 15 | 16 | 5 |
| 57 | 16 | 50 | 5 |
| 57 | 16 | 5 | 5 |
| 57 | 16 | 15 | 5 |
| 50 | 16 | 5 | 5 |
| 50 | 16 | 15 | 5 |
| 5 | 16 | 15 | 5 |

| | | | |
|---|---|---|---|
| 18 | 17 | 51 | 5 |
| 18 | 17 | 52 | 5 |
| 18 | 17 | 58 | 5 |
| 51 | 17 | 52 | 5 |
| 51 | 17 | 58 | 5 |
| 52 | 17 | 58 | 5 |
| 17 | 18 | 2 | 5 |
| 17 | 18 | 19 | 5 |
| 17 | 18 | 53 | 5 |
| 2 | 18 | 19 | 5 |
| 2 | 18 | 53 | 5 |
| 19 | 18 | 53 | 5 |
| 25 | 19 | 18 | 5 |
| 25 | 19 | 20 | 5 |
| 18 | 19 | 20 | 5 |
| 19 | 20 | 21 | 5 |
| 19 | 20 | 54 | 5 |
| 21 | 20 | 54 | 5 |
| 20 | 21 | 22 | 5 |
| 20 | 21 | 55 | 5 |
| 22 | 21 | 55 | 5 |
| 21 | 22 | 23 | 5 |
| 21 | 22 | 24 | 5 |
| 23 | 22 | 24 | 5 |
| 25 | 24 | 22 | 5 |
| 19 | 25 | 56 | 5 |
| 19 | 25 | 24 | 5 |
| 56 | 25 | 24 | 5 |
| 17 | 52 | 61 | 5 |
| 2 | 57 | 59 | 5 |
| 2 | 57 | 58 | 5 |
| 2 | 57 | 16 | 5 |
| 59 | 57 | 58 | 5 |
| 59 | 57 | 16 | 5 |
| 58 | 57 | 16 | 5 |
| 17 | 58 | 57 | 5 |
| 17 | 58 | 59 | 5 |
| 17 | 58 | 60 | 5 |
| 57 | 58 | 59 | 5 |
| 57 | 58 | 60 | 5 |
| 59 | 58 | 60 | 5 |
| 57 | 59 | 58 | 5 |
| 52 | 61 | 62 | 5 |
| 52 | 61 | 63 | 5 |
| 62 | 61 | 63 | 5 |
| 65 | 62 | 66 | 5 |

```
   65    62    61   5
   65    62    64   5
   66    62    61   5
   66    62    64   5
   61    62    64   5

[ dihedrals ]
; ai   aj ak  al funct  c0 c1  c2 c3   c4 c5
   27     1     2    18                    9
   27     1     2     3                    9
   27     1     2    57                    9
   28     1     2    18                    9
   28     1     2     3                    9
   28     1     2    57                    9
   26     1     2    18                    9
   26     1     2     3                    9
   26     1     2    57                    9
    1     2    18    17                    9
    1     2    18    19                    9
    1     2    18    53                    9
    3     2    18    17                    9
    3     2    18    19                    9
    3     2    18    53                    9
   57     2    18    17                    9
   57     2    18    19                    9
   57     2    18    53                    9
    1     2     3     4                    9
    1     2     3    29                    9
    1     2     3    30                    9
   18     2     3     4                    9
   18     2     3    29                    9
   18     2     3    30                    9
   57     2     3     4                    9
   57     2     3    29                    9
   57     2     3    30                    9
    1     2    57    59                    9
    1     2    57    58                    9
    1     2    57    16                    9
   18     2    57    59                    9
   18     2    57    58                    9
   18     2    57    16                    9
    3     2    57    59                    9
    3     2    57    58                    9
    3     2    57    16                    9
    2     3     4     5                    9
    2     3     4    31                    9
```

| | | | | |
|---|---|---|---|---|
| 2 | 3 | 4 | 32 | 9 |
| 29 | 3 | 4 | 5 | 9 |
| 29 | 3 | 4 | 31 | 9 |
| 29 | 3 | 4 | 32 | 9 |
| 30 | 3 | 4 | 5 | 9 |
| 30 | 3 | 4 | 31 | 9 |
| 30 | 3 | 4 | 32 | 9 |
| 3 | 4 | 5 | 33 | 9 |
| 3 | 4 | 5 | 6 | 9 |
| 3 | 4 | 5 | 16 | 9 |
| 31 | 4 | 5 | 33 | 9 |
| 31 | 4 | 5 | 6 | 9 |
| 31 | 4 | 5 | 16 | 9 |
| 32 | 4 | 5 | 33 | 9 |
| 32 | 4 | 5 | 6 | 9 |
| 32 | 4 | 5 | 16 | 9 |
| 33 | 5 | 6 | 13 | 9 |
| 33 | 5 | 6 | 7 | 9 |
| 33 | 5 | 6 | 8 | 9 |
| 4 | 5 | 6 | 13 | 9 |
| 4 | 5 | 6 | 7 | 9 |
| 4 | 5 | 6 | 8 | 9 |
| 16 | 5 | 6 | 13 | 9 |
| 16 | 5 | 6 | 7 | 9 |
| 16 | 5 | 6 | 8 | 9 |
| 33 | 5 | 16 | 57 | 9 |
| 33 | 5 | 16 | 50 | 9 |
| 33 | 5 | 16 | 15 | 9 |
| 4 | 5 | 16 | 57 | 9 |
| 4 | 5 | 16 | 50 | 9 |
| 4 | 5 | 16 | 15 | 9 |
| 6 | 5 | 16 | 57 | 9 |
| 6 | 5 | 16 | 50 | 9 |
| 6 | 5 | 16 | 15 | 9 |
| 5 | 6 | 13 | 12 | 9 |
| 5 | 6 | 13 | 45 | 9 |
| 5 | 6 | 13 | 14 | 9 |
| 7 | 6 | 13 | 12 | 9 |
| 7 | 6 | 13 | 45 | 9 |
| 7 | 6 | 13 | 14 | 9 |
| 8 | 6 | 13 | 12 | 9 |
| 8 | 6 | 13 | 45 | 9 |
| 8 | 6 | 13 | 14 | 9 |
| 13 | 6 | 7 | 34 | 9 |
| 13 | 6 | 7 | 35 | 9 |
| 13 | 6 | 7 | 36 | 9 |

| | | | | |
|---|---|---|---|---|
| 5 | 6 | 7 | 34 | 9 |
| 5 | 6 | 7 | 35 | 9 |
| 5 | 6 | 7 | 36 | 9 |
| 8 | 6 | 7 | 34 | 9 |
| 8 | 6 | 7 | 35 | 9 |
| 8 | 6 | 7 | 36 | 9 |
| 13 | 6 | 8 | 9 | 9 |
| 13 | 6 | 8 | 37 | 9 |
| 13 | 6 | 8 | 38 | 9 |
| 5 | 6 | 8 | 9 | 9 |
| 5 | 6 | 8 | 37 | 9 |
| 5 | 6 | 8 | 38 | 9 |
| 7 | 6 | 8 | 9 | 9 |
| 7 | 6 | 8 | 37 | 9 |
| 7 | 6 | 8 | 38 | 9 |
| 37 | 8 | 9 | 10 | 9 |
| 37 | 8 | 9 | 40 | 9 |
| 37 | 8 | 9 | 39 | 9 |
| 6 | 8 | 9 | 10 | 9 |
| 6 | 8 | 9 | 40 | 9 |
| 6 | 8 | 9 | 39 | 9 |
| 38 | 8 | 9 | 10 | 9 |
| 38 | 8 | 9 | 40 | 9 |
| 38 | 8 | 9 | 39 | 9 |
| 40 | 9 | 10 | 41 | 9 |
| 40 | 9 | 10 | 11 | 9 |
| 40 | 9 | 10 | 12 | 9 |
| 39 | 9 | 10 | 41 | 9 |
| 39 | 9 | 10 | 11 | 9 |
| 39 | 9 | 10 | 12 | 9 |
| 8 | 9 | 10 | 41 | 9 |
| 8 | 9 | 10 | 11 | 9 |
| 8 | 9 | 10 | 12 | 9 |
| 9 | 10 | 11 | 42 | 9 |
| 41 | 10 | 11 | 42 | 9 |
| 12 | 10 | 11 | 42 | 9 |
| 9 | 10 | 12 | 43 | 9 |
| 9 | 10 | 12 | 44 | 9 |
| 9 | 10 | 12 | 13 | 9 |
| 41 | 10 | 12 | 43 | 9 |
| 41 | 10 | 12 | 44 | 9 |
| 41 | 10 | 12 | 13 | 9 |
| 11 | 10 | 12 | 43 | 9 |
| 11 | 10 | 12 | 44 | 9 |
| 11 | 10 | 12 | 13 | 9 |
| 10 | 12 | 13 | 45 | 9 |

| | | | | |
|---|---|---|---|---|
| 10 | 12 | 13 | 6 | 9 |
| 10 | 12 | 13 | 14 | 9 |
| 43 | 12 | 13 | 45 | 9 |
| 43 | 12 | 13 | 6 | 9 |
| 43 | 12 | 13 | 14 | 9 |
| 44 | 12 | 13 | 45 | 9 |
| 44 | 12 | 13 | 6 | 9 |
| 44 | 12 | 13 | 14 | 9 |
| 12 | 13 | 14 | 47 | 9 |
| 12 | 13 | 14 | 46 | 9 |
| 12 | 13 | 14 | 15 | 9 |
| 45 | 13 | 14 | 47 | 9 |
| 45 | 13 | 14 | 46 | 9 |
| 45 | 13 | 14 | 15 | 9 |
| 6 | 13 | 14 | 47 | 9 |
| 6 | 13 | 14 | 46 | 9 |
| 6 | 13 | 14 | 15 | 9 |
| 47 | 14 | 15 | 49 | 9 |
| 47 | 14 | 15 | 48 | 9 |
| 47 | 14 | 15 | 16 | 9 |
| 13 | 14 | 15 | 49 | 9 |
| 13 | 14 | 15 | 48 | 9 |
| 13 | 14 | 15 | 16 | 9 |
| 46 | 14 | 15 | 49 | 9 |
| 46 | 14 | 15 | 48 | 9 |
| 46 | 14 | 15 | 16 | 9 |
| 49 | 15 | 16 | 57 | 9 |
| 49 | 15 | 16 | 50 | 9 |
| 49 | 15 | 16 | 5 | 9 |
| 48 | 15 | 16 | 57 | 9 |
| 48 | 15 | 16 | 50 | 9 |
| 48 | 15 | 16 | 5 | 9 |
| 14 | 15 | 16 | 57 | 9 |
| 14 | 15 | 16 | 50 | 9 |
| 14 | 15 | 16 | 5 | 9 |
| 50 | 16 | 57 | 2 | 9 |
| 50 | 16 | 57 | 59 | 9 |
| 50 | 16 | 57 | 58 | 9 |
| 5 | 16 | 57 | 2 | 9 |
| 5 | 16 | 57 | 59 | 9 |
| 5 | 16 | 57 | 58 | 9 |
| 15 | 16 | 57 | 2 | 9 |
| 15 | 16 | 57 | 59 | 9 |
| 15 | 16 | 57 | 58 | 9 |
| 51 | 17 | 18 | 2 | 9 |
| 51 | 17 | 18 | 19 | 9 |

| | | | | |
|---|---|---|---|---|
| 51 | 17 | 18 | 53 | 9 |
| 52 | 17 | 18 | 2 | 9 |
| 52 | 17 | 18 | 19 | 9 |
| 52 | 17 | 18 | 53 | 9 |
| 58 | 17 | 18 | 2 | 9 |
| 58 | 17 | 18 | 19 | 9 |
| 58 | 17 | 18 | 53 | 9 |
| 18 | 17 | 52 | 61 | 9 |
| 51 | 17 | 52 | 61 | 9 |
| 58 | 17 | 52 | 61 | 9 |
| 18 | 17 | 58 | 57 | 9 |
| 18 | 17 | 58 | 59 | 9 |
| 18 | 17 | 58 | 60 | 9 |
| 51 | 17 | 58 | 57 | 9 |
| 51 | 17 | 58 | 59 | 9 |
| 51 | 17 | 58 | 60 | 9 |
| 52 | 17 | 58 | 57 | 9 |
| 52 | 17 | 58 | 59 | 9 |
| 52 | 17 | 58 | 60 | 9 |
| 17 | 18 | 19 | 25 | 9 |
| 17 | 18 | 19 | 20 | 9 |
| 2 | 18 | 19 | 25 | 9 |
| 2 | 18 | 19 | 20 | 9 |
| 53 | 18 | 19 | 25 | 9 |
| 53 | 18 | 19 | 20 | 9 |
| 18 | 19 | 25 | 56 | 9 |
| 18 | 19 | 25 | 24 | 9 |
| 20 | 19 | 25 | 56 | 9 |
| 20 | 19 | 25 | 24 | 9 |
| 25 | 19 | 20 | 21 | 9 |
| 25 | 19 | 20 | 54 | 9 |
| 18 | 19 | 20 | 21 | 9 |
| 18 | 19 | 20 | 54 | 9 |
| 19 | 20 | 21 | 22 | 9 |
| 19 | 20 | 21 | 55 | 9 |
| 54 | 20 | 21 | 22 | 9 |
| 54 | 20 | 21 | 55 | 9 |
| 20 | 21 | 22 | 23 | 9 |
| 20 | 21 | 22 | 24 | 9 |
| 55 | 21 | 22 | 23 | 9 |
| 55 | 21 | 22 | 24 | 9 |
| 21 | 22 | 24 | 25 | 9 |
| 23 | 22 | 24 | 25 | 9 |
| 22 | 24 | 25 | 19 | 9 |
| 22 | 24 | 25 | 56 | 9 |
| 17 | 52 | 61 | 62 | 9 |

```
   17    52    61    63                      9
    2    57    59    58                      9
   16    57    59    58                      9
    2    57    58    17                      9
    2    57    58    59                      9
    2    57    58    60                      9
   59    57    58    17                      9
   59    57    58    60                      9
   16    57    58    17                      9
   16    57    58    59                      9
   16    57    58    60                      9
   17    58    59    57                      9
   60    58    59    57                      9
   52    61    62    65                      9
   52    61    62    66                      9
   52    61    62    64                      9
   63    61    62    65                      9
   63    61    62    66                      9
   63    61    62    64                      9

[ dihedrals ]
; ai  aj ak  al funct  c0 c1  c2 c3
   22    21    23    24                      2
   61    62    63    52                      2
```

**Supporting data file S4.** Cinobufagin prm file.

```
[ bondtypes ]
;      i        j  func           b0           kb
  CG2R62   CG3C51     1   0.14900000    192464.00
  CG3C51    OG302     1   0.14110000    279742.24

[ angletypes ]
;      i        j        k  func       theta0       ktheta          ub0          kub
  CG2R62   CG2R62   CG3C51     5   124.200000   334.720000   0.00000000         0.00
  OG3R60   CG2R62    HGR62     5   119.000000   292.880000   0.00000000         0.00
  CG2R62   CG3C51   CG3C51     5   112.300000   435.136000   0.00000000         0.00
  CG2R62   CG3C51   CG3RC1     5   112.300000   435.136000   0.00000000         0.00
  CG2R62   CG3C51     HGA1     5   112.000000   418.400000   0.00000000         0.00
  CG3C51   CG3C51    OG302     5   106.500000   485.344000   0.25610000      6694.40
  CG3RC1   CG3C51    OG302     5   110.100000   633.457600   0.00000000         0.00
   OG302   CG3C51     HGA1     5   108.500000   384.091200   0.00000000         0.00
   CG311   CG3RC1   OG3C31     5   115.500000   505.008800   0.00000000         0.00
  CG3RC1   CG3RC1   CG3RC1     5   109.000000   418.400000   0.00000000         0.00
   CG2O2    OG302   CG3C51     5   109.600000   334.720000   0.22651000     25104.00

[ dihedraltypes ]
;      i        j        k        l  func         phi0         kphi  mult
   CG331    CG2O2    OG302   CG3C51     9   180.000000     8.577200     2
   OG2D1    CG2O2    OG302   CG3C51     9   180.000000     4.037560     1
   OG2D1    CG2O2    OG302   CG3C51     9   180.000000    16.108400     2
  CG2R62   CG2R62   CG2R62   CG3C51     9   180.000000    12.970400     2
  CG3C51   CG2R62   CG2R62   OG3R60     9   180.000000    10.041600     2
  CG3C51   CG2R62   CG2R62    HGR62     9   180.000000    16.736000     2
  CG2R62   CG2R62   CG3C51   CG3C51     9   180.000000     0.962320     2
  CG2R62   CG2R62   CG3C51   CG3RC1     9   180.000000     0.962320     2
  CG2R62   CG2R62   CG3C51     HGA1     9     0.000000     0.008368     6
   HGR62   CG2R62   OG3R60   CG2R63     9     0.000000     3.179840     2
   CG311    CG311   CG3RC1   OG3C31     9   180.000000     1.966480     1
   CG311    CG311   CG3RC1   OG3C31     9     0.000000     0.543920     3
   CG321    CG311   CG3RC1   OG3C31     9   180.000000     1.966480     1
   CG321    CG311   CG3RC1   OG3C31     9     0.000000     0.543920     3
    HGA1    CG311   CG3RC1   OG3C31     9     0.000000     0.669440     3
  CG2R62   CG3C51   CG3C51   CG3RC1     9     0.000000     0.585760     3
  CG2R62   CG3C51   CG3C51    OG302     9     0.000000     0.000000     3
  CG2R62   CG3C51   CG3C51     HGA1     9     0.000000     0.585760     3
  CG3RC1   CG3C51   CG3C51   CG3RC1     9     0.000000     0.627600     3
  CG3RC1   CG3C51   CG3C51    OG302     9   180.000000     3.347200     3
  CG3RC1   CG3C51   CG3C51    OG302     9     0.000000     0.836800     4
   OG302   CG3C51   CG3C51     HGA1     9     0.000000     0.815880     3
  CG2R62   CG3C51   CG3RC1    CG321     9   180.000000     2.092000     2
  CG2R62   CG3C51   CG3RC1    CG331     9   180.000000     2.092000     2
```

| | | | | | | | |
|---|---|---|---|---|---|---|---|
| CG2R62 | CG3C51 | CG3RC1 | CG3RC1 | 9 | 0.000000 | 0.627600 | 3 |
| CG3C51 | CG3C51 | CG3RC1 | CG321 | 9 | 180.000000 | 9.204800 | 2 |
| CG3C51 | CG3C51 | CG3RC1 | CG321 | 9 | 0.000000 | 16.736000 | 3 |
| CG3C51 | CG3C51 | CG3RC1 | CG321 | 9 | 180.000000 | 2.301200 | 6 |
| CG3C51 | CG3C51 | CG3RC1 | CG331 | 9 | 0.000000 | 0.209200 | 3 |
| OG302 | CG3C51 | CG3RC1 | CG3RC1 | 9 | 0.000000 | 1.882800 | 2 |
| OG302 | CG3C51 | CG3RC1 | CG3RC1 | 9 | 0.000000 | 3.765600 | 6 |
| OG302 | CG3C51 | CG3RC1 | OG3C31 | 9 | 0.000000 | 0.836800 | 3 |
| OG302 | CG3C51 | CG3RC1 | OG3C31 | 9 | 180.000000 | 2.510400 | 4 |
| OG302 | CG3C51 | CG3RC1 | OG3C31 | 9 | 0.000000 | 1.255200 | 5 |
| OG302 | CG3C51 | CG3RC1 | OG3C31 | 9 | 0.000000 | 2.092000 | 6 |
| OG302 | CG3C51 | CG3RC1 | HGA1 | 9 | 0.000000 | 0.627600 | 3 |
| CG3C51 | CG3C51 | OG302 | CG2O2 | 9 | 180.000000 | 0.418400 | 1 |
| CG3C51 | CG3C51 | OG302 | CG2O2 | 9 | 180.000000 | 6.903600 | 2 |
| CG3C51 | CG3C51 | OG302 | CG2O2 | 9 | 0.000000 | 1.882800 | 3 |
| CG3RC1 | CG3C51 | OG302 | CG2O2 | 9 | 180.000000 | 0.418400 | 1 |
| CG3RC1 | CG3C51 | OG302 | CG2O2 | 9 | 180.000000 | 6.903600 | 2 |
| CG3RC1 | CG3C51 | OG302 | CG2O2 | 9 | 0.000000 | 1.882800 | 3 |
| HGA1 | CG3C51 | OG302 | CG2O2 | 9 | 0.000000 | 0.000000 | 3 |
| CG311 | CG3RC1 | CG3RC1 | OG3C31 | 9 | 0.000000 | 0.627600 | 3 |
| CG321 | CG3RC1 | CG3RC1 | CG3RC1 | 9 | 0.000000 | 0.627600 | 3 |
| CG321 | CG3RC1 | CG3RC1 | OG3C31 | 9 | 0.000000 | 0.627600 | 3 |
| CG331 | CG3RC1 | CG3RC1 | CG3RC1 | 9 | 0.000000 | 0.209200 | 3 |
| CG331 | CG3RC1 | CG3RC1 | OG3C31 | 9 | 0.000000 | 0.627600 | 3 |
| CG3C51 | CG3RC1 | CG3RC1 | CG3RC1 | 9 | 0.000000 | 16.736000 | 3 |
| CG3RC1 | CG3RC1 | CG3RC1 | OG3C31 | 9 | 0.000000 | 5.020800 | 3 |
| CG3RC1 | CG3RC1 | CG3RC1 | HGA1 | 9 | 0.000000 | 0.627600 | 3 |
| CG311 | CG3RC1 | OG3C31 | CG3RC1 | 9 | 0.000000 | 2.677760 | 5 |
| CG311 | CG3RC1 | OG3C31 | CG3RC1 | 9 | 180.000000 | 3.389040 | 6 |
| CG3RC1 | CG3RC1 | OG3C31 | CG3RC1 | 9 | 0.000000 | 2.677760 | 5 |
| CG3RC1 | CG3RC1 | OG3C31 | CG3RC1 | 9 | 180.000000 | 3.389040 | 6 |

[ dihedraltypes ]
; 'improper' dihedrals
; i j k l func phi0 kphi
